# Towards a gold standard for benchmarking gene set enrichment analysis

**DOI:** 10.1101/674267

**Authors:** Ludwig Geistlinger, Gergely Csaba, Mara Santarelli, Marcel Ramos, Lucas Schiffer, Charity Law, Nitesh Turaga, Sean Davis, Vincent Carey, Martin Morgan, Ralf Zimmer, Levi Waldron

## Abstract

**Background:** Although gene set enrichment analysis has become an integral part of high-throughput gene expression data analysis, the assessment of enrichment methods remains rudimentary and ad hoc. In the absence of suitable gold standards, evaluations are commonly restricted to selected data sets and biological reasoning on the relevance of resulting enriched gene sets. However, this is typically incomplete and biased towards the goals of individual investigations.

**Results:** We present a general framework for standardized and structured benchmarking of enrichment methods based on defined criteria for applicability, gene set prioritization, and detection of relevant processes. This framework incorporates a curated compendium of 75 expression data sets investigating 42 different human diseases. The compendium features microarray and RNA-seq measurements, and each dataset is associated with a precompiled GO/KEGG relevance ranking for the corresponding disease under investigation. We perform a comprehensive assessment of 10 major enrichment methods on the benchmark compendium, identifying significant differences in (i) runtime and applicability to RNA-seq data, (ii) fraction of enriched gene sets depending on the type of null hypothesis tested, and (iii) recovery of the *a priori* defined relevance rankings. Based on these findings, we make practical recommendations on (i) how methods originally developed for microarray data can efficiently be applied to RNA-seq data, (ii) how to interpret results depending on the type of gene set test conducted, and (iii) which methods are best suited to effectively prioritize gene sets with high relevance for the phenotype investigated.

**Conclusion:** We carried out a systematic assessment of existing enrichment methods, and identified best performing methods, but also general shortcomings in how gene set analysis is currently conducted. We provide a directly executable benchmark system for straightforward assessment of additional enrichment methods.

**Availability:** http://bioconductor.org/packages/GSEABenchmarkeR

## Introduction

The goal of genome-wide gene expression studies is to discover the molecular mechanisms that underlie certain phenotypes such as human diseases [1]. For this purpose, expression changes of individual genes are typically analyzed for enrichment in functional gene sets. These sets represent molecular functions and biological processes as defined by the Gene Ontology [GO, 2] or the KEGG pathway database [3].

The two predominantly used enrichment methods are (i) overrepresentation analysis [ORA, 4], testing whether a gene set contains disproportionately many genes of significant expression change; and (ii) gene set enrichment analysis [GSEA, 5], rather testing whether genes of a gene set accumulate at the top or bottom of the full gene vector ordered by direction and magnitude of expression change.

However, the term *gene set enrichment analysis* now encompasses a general strategy implemented by a wide range of methods [6]. Those methods share a common goal, although approach and statistical model vary substantially [4, 7].

A major question is thus which method is best suited for the enrichment analysis. As a consequence, many methods have been published claiming improvement especially with respect to ORA and the original GSEA method. This claim is typically made based on (i) simulated data, specifically designed to demonstrate beneficial aspects of a new method, and (ii) real datasets, for which however the truly enriched gene sets are not known.

As the evaluation is thus typically based on self-defined standards including only a few methods, Mitrea *et al*. [8] identified the lack of gold standards for consistent assessment and comparison of enrichment methods as a major bottleneck. Steps towards an objective assessment are recent independent studies [9–13], which evaluated a partly overlapping selection of enrichment methods on (i) simulated data, modeling certain aspects of real data [9]; (ii) real data, arguing on the biological relevance of the enriched gene sets [10]; or (iii) a combination of both data types [11–14].

As the standard of truth is hard to establish for real data, several approaches have been suggested to *a priori* define target gene sets for specific datasets. For example, Naeem *et al*. [15] suggested an assessment based on known target gene sets of transcription factors for expression datasets where those transcription factors are overexpressed or knocked out as available for *E*. *coli* and *S*. *cerevisiae*. On the other hand, Tarca *et al*. [16, 17] collected 42 microarray datasets investigating human diseases for which a specific KEGG target pathway exists. This strategy has been adapted by several recent enrichment evaluation studies [18–21].

However, there is little agreement among studies on which methods to prefer, with most studies concluding with a recommendation for a consensus/combination of methods [12, 14, 15, 20]. Although this is valuable in practice, existing assessments (i) were mostly based on microarray data, and it is not clear whether results hold equally for RNA-seq data, (ii) do not represent the wide range of existing methods, and (iii) are often cumbersome to reproduce for additional methods, as this involves considerable effort of data processing and method collection.

## Results

We present the `GSEABenchmarkeR` R/Bioconductor package, which implements an executable benchmark framework for the systematic and reproducible assessment of gene set enrichment methods (Figure 1). The package facilitates efficient execution of a representative and extendable collection of enrichment analysis (EA) methods on comprehensive real data compendia. The compendia are curated collections of microarray and RNA-seq datasets investigating human diseases (mostly specific cancer types), for which disease-relevant gene sets have been defined *a priori*. Consistently applied to these datasets, methods can then be assessed with respect to computational runtime, statistical significance, and phenotype relevance, i.e. whether methods produce gene set rankings in which phenotype-relevant gene sets accumulate at the top.

**Figure 1.**
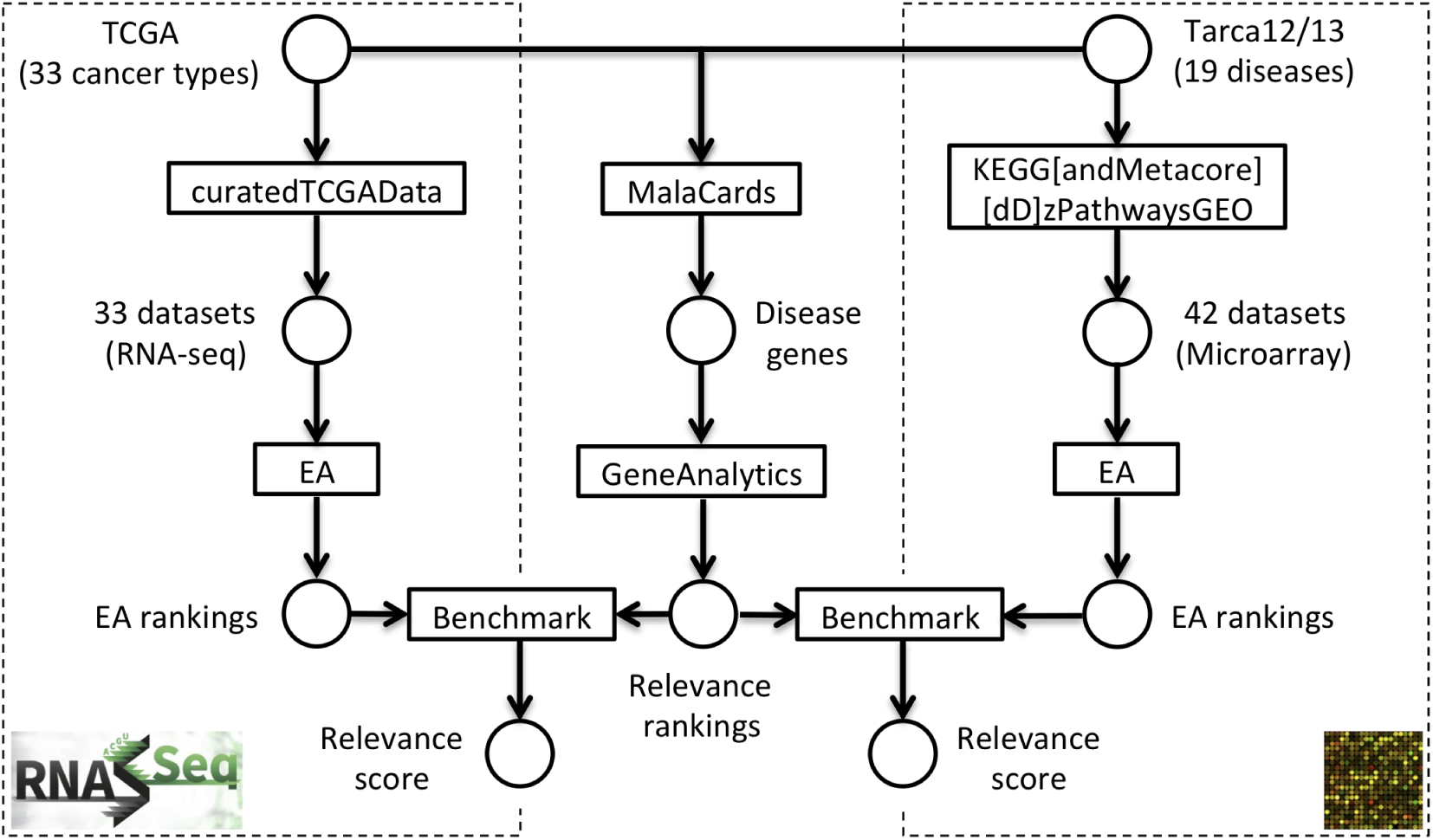
Benchmark setup. The benchmark framework incorporates a pre-defined RNA-seq panel (left), gene set relevance rankings (center), and a microarray panel (right). The RNA-seq panel investigates 33 cancer types across 33 datasets from TCGA [24], which are accessed through the curatedTCGAData package. The microarray panel investigates 19 human diseases across 42 datasets collected by Tarca *et al*. [16, 17], and which are available in the KEGGdzPathwaysGEO and KEGGandMetacoreDzPathwaysGEO package. Gene set relevance rankings for both data panels are constructed by (i) querying the MalaCards database [38] for each disease investigated, and (ii) subjecting resulting disease genes to GeneAnalytics [52], which yields relevance rankings for GO-BP terms and KEGG pathways. Enrichment analysis (EA) methods selected for benchmarking are carried out across datasets of the data panels, yielding a gene set ranking (EA ranking) for each method on each dataset. The resulting EA rankings for each dataset are then benchmarked against the precompiled relevance rankings for the corresponding disease investigated.

In the following, we use the package to assess the performance of 10 major enrichment analysis (EA) methods listed in Table 1. These methods represent a decade of developments and are well-established as indicated by their citation frequency. Following previously introduced nomenclature [20], we note that these are *set-based* methods, and thus ignore known interactions between genes. We also note that benchmarking with the `GSEABenchmarkeR` package extends to *network-based* methods that incorporate known interactions (Supplementary Methods S1.1). However, as the assessment of network-based methods require to additionally evaluate the choice of the network [22, 23], we decided to deal with these methods in a separate manuscript.

**Table 1.**
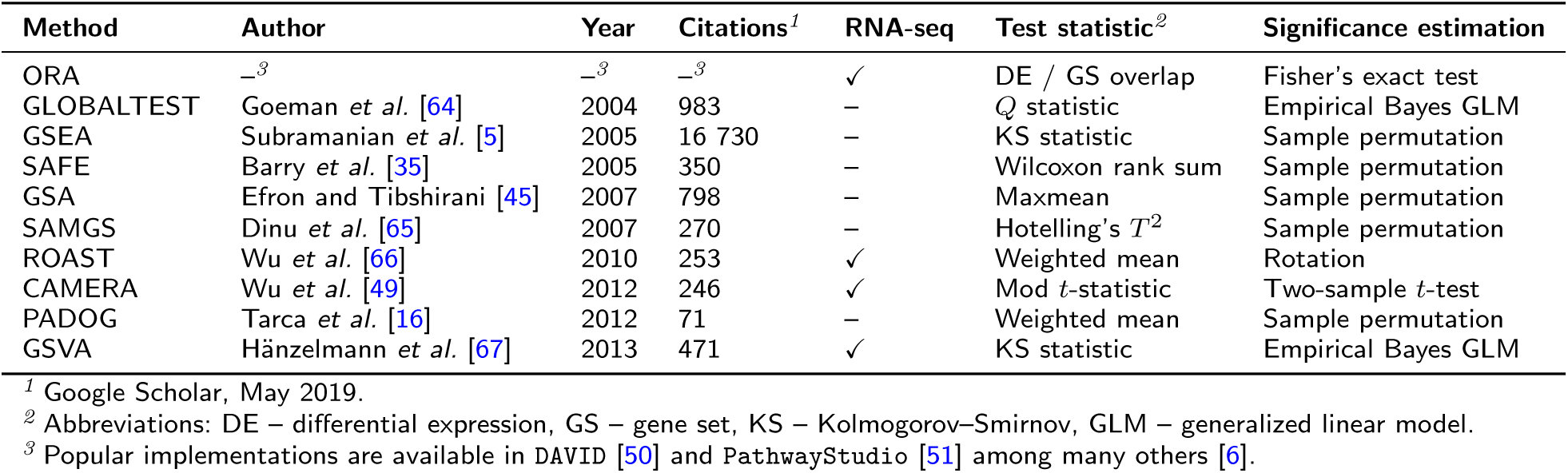
Gene set analysis methods under benchmark.

We start by exploring the benchmark compendia for sample size and differential expression (DE), and subsequently describe how EA methods developed for microarray data can be adapted for application to RNA-seq data.

### Benchmark compendia and gene set collections

As illustrated in Figure 1, the two pre-defined benchmark compendia consist of 42 microarray datasets collected by Tarca *et al*. [16, 17] and 33 RNA-seq datasets from The Cancer Genome Atlas [TCGA, 24]. These datasets investigate 42 human diseases, including 35 cancer types (Supplementary Tables S1 and S2).

When analyzing datasets of the benchmark compendia for sample size and differential expression (DE), we find them to display a representative range (Supplementary Figure S1). The 42 datasets of the GEO2KEGG microarray compendium range from a minimum of 4 cases and 4 controls to a maximum of 91 cases and 62 controls. Using the typical DE thresholds of (i) absolute log2 fold change > 1, and (ii) Benjamini-Hochberg [BH, 25]-adjusted *p*-value *<* 0.05, we find several datasets of the GEO2KEGG microarray compendium with not a single DE gene, and at the other extreme, datasets with up to 73% DE genes (according to the *p*-value criterion; up to 15% satisfying both criteria).

For this study, we restrict analysis of the TCGA RNA-seq compendium to cancer types for which at least 5 adjacent normal tissue samples are available and take the pairing of samples (tumor vs. adjacent normal) into account. This yields 15 cancer types / datasets ranging from a minimum of 9 patients to a maximum of 226 patients, for which both tumor and adjacent normal samples were available (Supplementary Figure S1). Datasets of the TCGA RNA-seq compendium display relatively high levels of differential expression, with a range of 34%-79% DE genes (according to the *p*-value criterion; 9%-29% satisfying both criteria).

When exploring the gene set size distribution in human KEGG pathways and GO-BP terms (Supplementary Figure S2), restricted to gene sets with a minimum of 5 genes and a maximum of 500 genes (the typical EA thresholds), we find that (i) there are considerable more GO-BP sets than KEGG sets (4,631 vs. 323), and (ii) GO-BP gene sets tend to be smaller (median set size: 11 vs. 72).

### Dealing with RNA-seq data

There is disagreement over whether EA methods originally developed for microarray data are directly applicable to RNA-seq data. This is partly caused by the variety of RNA-seq expression units that are used at different steps of RNA-seq data analysis. For instance, popular methods for DE analysis require the raw RNA-seq read counts as input to preserve the sampling characteristics of the data [26–28]. However, frequently used tools for transcript abundance estimation report transcripts per million [TPMs, 29] or fragments / reads per kilobase of transcript per million mapped reads [FPKMs/RPKMs, 30] that already account for differences in gene length and sequencing depth. As FPKM/RPKM is inconsistent between samples and can be directly converted to TPM [29, 31], we consider raw read counts or TPMs as input for the EA methods under benchmark.

Due to the different statistical models and implementations of the EA methods (Table 1), it is necessary to distinguish between *(i)* methods that work on the list of DE genes (ORA), which can be straightforward applied assuming that gene length bias is controlled for [32], *(ii)* methods that distinguish between a microarray mode, and an RNA-seq mode that assumes that the raw read counts are provided (CAMERA, ROAST, and GSVA), or *(iii)* methods that incorporate sample permutation and recalculation of *t*-like statistics for each gene (GSEA, SAFE, GSA, SAMGS, PADOG).

Methods of the third type require either a variance-stabilizing transformation [VST, 26, 33], or incorporation of RNA-seq tools such as `voom`/`limma`, `edgeR`, or `DESeq2` for calculation of the per-gene statistic in each permutation [20, 34]. Incorporation of RNA-seq tools is straightforward for the permutation framework implemented in SAFE [35] as it allows to provide user-defined local (per-gene) and global (gene set) test statistics.

For the following assessment of EA methods, we thus also analyze the differences of using raw counts or VST-transformed counts as input. However, for the datasets of TCGA RNA-seq compendium, we observe almost identical fold changes and DE *p*-values when using either (1) `voom`/`limma` on raw read counts or TPMs, or (2) `limma` on VST-transformed counts or log TPMs (Supplementary Tables S3 and S4).

### Runtime

Computational runtime is an important measure of the applicability of a method. When analyzing enrichment methods on the microarray compendium using GO-BP gene sets (Figure 2), average runtime ranged from a minimum of 7.7 sec (CAMERA) to a maximum of 32.6 min (GSEA). Closer inspection reveals three groups of methods reflecting aspects of methodology and implementation (Table 1). CAMERA, ORA, and GLOBALTEST use simple parametric tests for gene set significance estimation, which results in fast runtimes. The other methods are computationally more intensive as they use sample permutation (SAFE, SAMGS, GSA, PADOG, and GSEA) or Monte Carlo sampling (GSVA and ROAST). The most computationally expensive are GSA (additional gene permutation), PADOG (gene weighting by occurrence frequency), and GSEA (cumulative KS-statistic). Runtimes on the TCGA RNA-seq compendium and when using KEGG gene sets displayed a similar pattern (Supplementary Figure S3). However, we observed significantly increased runtimes when carrying out methods with dedicated RNA-seq mode on raw read counts (Supplementary Figure S4). This is especially apparent for the case of incorporating RNA-seq tools in the SAFE framework, where runtime also depends on the RNA-seq tool used (`voom`/`limma ≪ edgeR ≪ DESeq2`).

**Figure 2.**
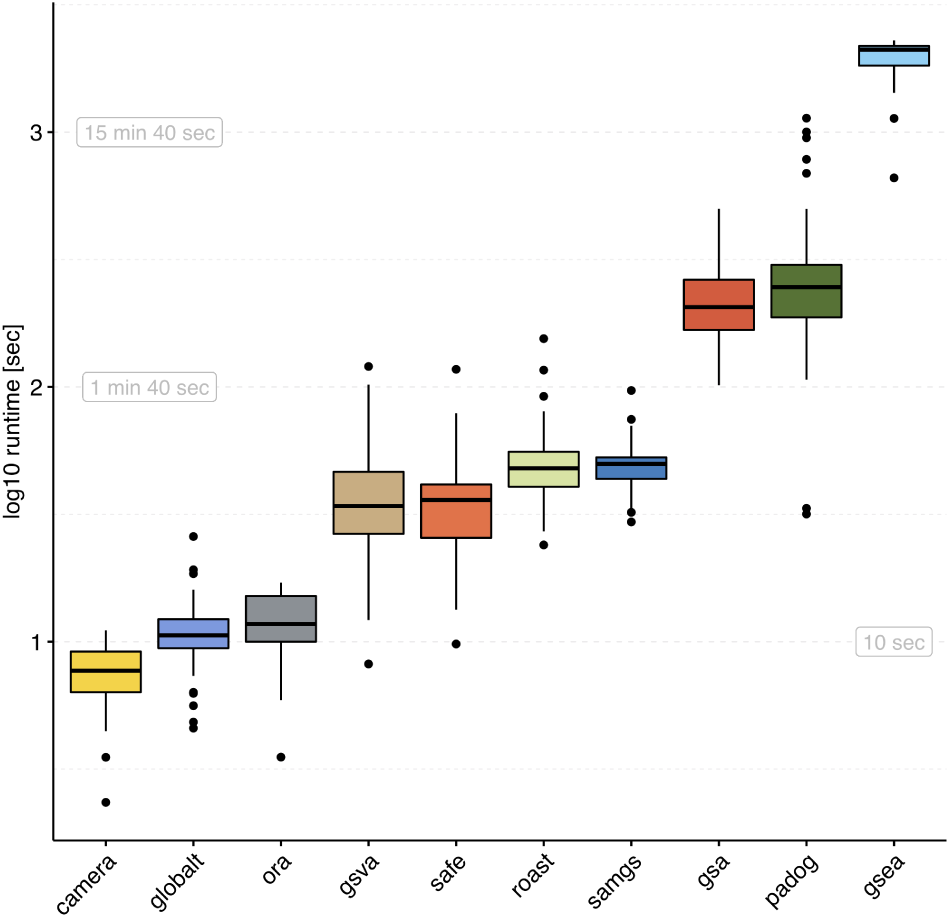
Runtime. Elapsed processing times (*y*-axis,log-scale) when applying the enrichment methods indicated on the *x*-axis to the 42 datasets of the GEO2KEGG microarray compendium. Gene sets were defined according to GO-BP (*N* = 4, 631). Computation was carried out on an Intel Xeon 2.7 GHz machine. Runtimes for the TCGA RNA-seq compendium and when using KEGG gene sets are shown in Supplementary Figure S3.

### Statistical significance

Enrichment methods conduct a hypothesis test for each gene set under investigation. The underlying null hypothesis can be characterized as either (i) *selfcontained* : no genes in the set of interest are DE, or (ii) *competitive*: the genes in the set of interest are at most as often DE as the genes not in the set [4]. As typically many gene sets are tested, multiple testing correction is needed to account for type I error rate inflation [36].

Using the popular BH-method [25] for multiple testing correction and an adjusted significance level of 0.05, we find EA methods to report drastically different fractions of gene sets as statistically significant (Figure 3). This is tied to the type of null hypothesis tested, with self-contained methods reporting much larger fractions of significant gene sets. Conversely, we find several competitive methods (SAFE, GSEA, GSA, PADOG) to frequently report not a single significant gene set, and two self-contained methods (GLOBALTEST and SAMGS) to frequently report all gene sets tested as significant.

**Figure 3.**
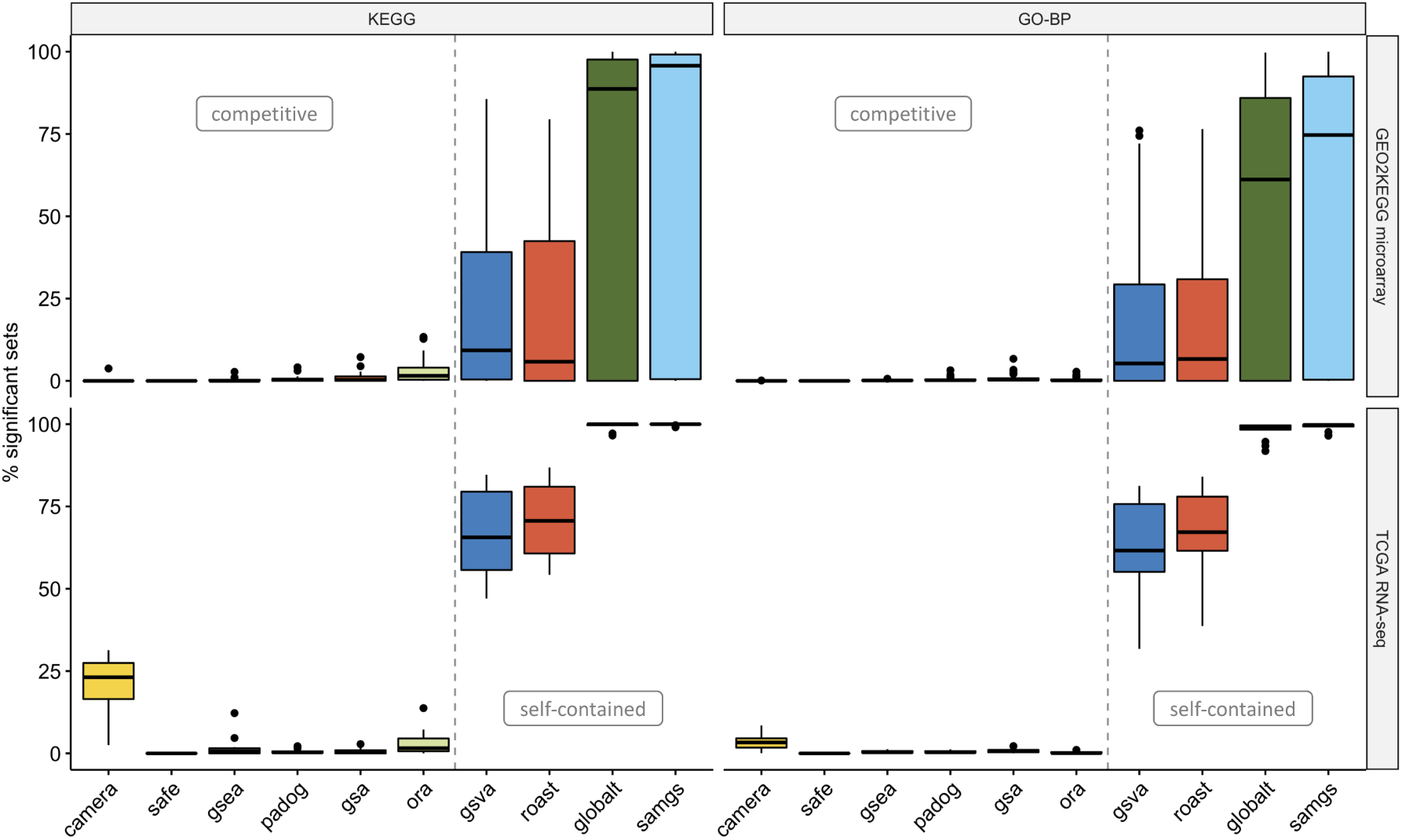
Statistical significance. Percentage of significant gene sets (FDR *<* 0.05, *y*-axis) when applying methods to the GEO2KEGG microarray compendium (top, 42 datasets) and the TCGA RNA-seq compendium (bottom, 15 datasets). Gene sets were defined according to KEGG (left, 323 gene sets) and GO-BP (right, 4,631 gene sets). The grey dashed line divides methods based on the type of null hypothesis tested [4]. Supplementary Figure S7 shows the percentage of significant gene sets when using a nominal significance threshold of 0.05.

To ensure correct application of methods, we applied them in a controlled setup (Figure 4). We therefore used the well-studied microarray dataset of Golub *et al*. [37] that contrasts the transcriptome profiles of acute myeloid leukemia (AML) and acute lymphoblastic leukemia (ALL) patients. By shuffling sample labels (AML vs. ALL) 1,000 times, and assessing in each permutation the number of GO-BP gene sets with *p <* 0.05, we find average type I error rates controlled at the 5%-level (Figure 4a). However, self-contained methods displayed in certain random assignments of the sample labels substantially elevated type I error rates. This effect was more pronounced for KEGG gene sets, which tend to be larger (Supplementary Figure S5). To test for a possible gene set size effect, we also applied methods to the Golub dataset (true sample labels) with randomly sampled gene sets of increasing size (Figure 4b). Self-contained methods reported systematically larger fractions of significant random gene sets, with GLOBALTEST and SAMGS displaying a set size dependency that resulted in rendering all random gene sets with >50 genes significant.

**Figure 4.**
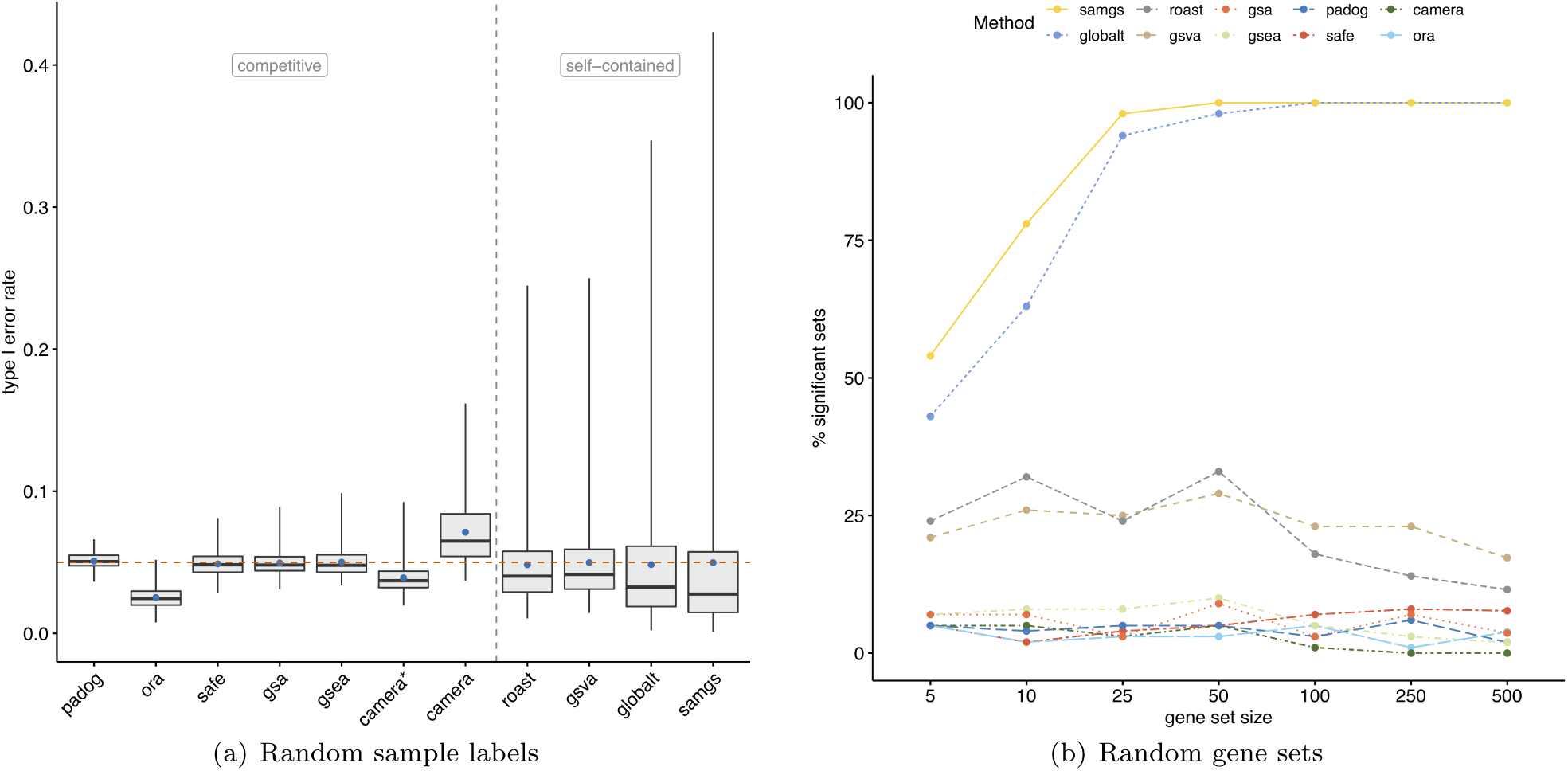
Random sample labels and random gene sets. **(a)** Type I error rates (*y*-axis) as evaluated on the Golub dataset [37] by shuffling sample labels 1000 times, and assessing in each permutation the fraction of gene sets with *p <* 0.05. Gene sets were defined according to GO-BP (*N* = 4, 631). Blue points indicate the mean type I error rate and the red dashed line the significance level of 0.05. The grey dashed line divides methods based on the type of null hypothesis tested [4]. Supplementary Figure S5 shows type I error rates when using KEGG gene sets. * application of CAMERA without accounting for inter-gene correlation (default: inter-gene correlation of 0.01). **(b)** Percentage of significant gene sets (*p <* 0.05, *y*-axis) when applying methods to the Golub dataset (true sample labels) and using 100 randomly sampled gene sets of defined size (*x*-axis).

This gene set size dependence was also apparent for both benchmark compendia, where self-contained methods reported larger fractions of significant gene sets for KEGG than for GO-BP (Figure 3). Following from the definition of the respective null hypothesis, self-contained but not competitive methods also display dependence on the background level of DE in a dataset (Supplementary Figure S6). As competitive methods were highly conservative, we inspected their nominal *p*-value distributions. Fraction of gene sets with nominal *p <* 0.05 were constant across datasets at ≈5-15% (Supplementary Figure S7), and the effect of the multiple testing correction was invariant to increasing the number of permutations or using the respective built-in FDR correction for GSEA and SAFE (Supplementary Figure S8).

### Phenotype relevance

Evaluations of published EA methods often conclude phenotype relevance if there is any association between the top-ranked gene sets and the investigated phenotype. This involves a certain extent of cherry-picking from the enriched gene sets, where sets with a link to the phenotype are preferentially selected. For an impartial assessment, we propose to rather investigate phenotype relevance of all gene sets *a priori*, and to subsequently quantify the relevance accumulated along the gene set ranking.

For the non-trivial task of scoring the phenotype relevance of a gene set, we build on the `MalaCards` disease database [38]. `MalaCards` scores genes for disease relevance based on experimental evidence and co-citation in the literature, and summarizes per-gene relevance across GO and KEGG gene sets. Focusing on the diseases investigated in the datasets of the benchmark compendia, we systematically extracted disease genes and gene set relevance rankings from `MalaCards` (see again Figure 1). As expected, disease genes and gene sets for cancer types studied in the benchmark compendia (Supplementary Figures S10 and S11) are enriched for known cancer driver genes and oncogenic processes [39, 40]. Relevance rankings are also more similar within disease classes than between disease classes (Supplementary Figure S13).

By scoring the similarity between the EA rankings and the precompiled relevance rankings, we assess whether certain EA methods tend to produce rankings of higher phenotype relevance (as outlined in Figure 1 and detailed in Methods, Section *Phenotype relevance*). We observed that competitive methods tend to rank phenotype-relevant gene sets systematically higher than self-contained methods (Figure 5). This observation holds for all 4 combinations of benchmark compendium (GEO2KEGG and TCGA) and gene set collection (KEGG and GO-BP), resulting in a significant overall difference between competitive and selfcontained methods (*p* = 1.87 · 10^-19^, Wilcoxon ranksum test). Differences between competitive methods were only moderate, with PADOG consistently returning highest relevance scores. However, PADOG scores were overall not significantly higher than for ORA (*p* = 0.85, Wilcoxon rank-sum test) and SAFE (*p* = 0.19), but significantly exceeded the scores of GSEA (*p* = 0.014), GSA (*p* = 0.04), and CAMERA (*p* = 0.002). We also confirmed that these observations largely hold when restricting the evaluation to the top 20% of each EA ranking (Supplementary Figure S14).

**Figure 5.**
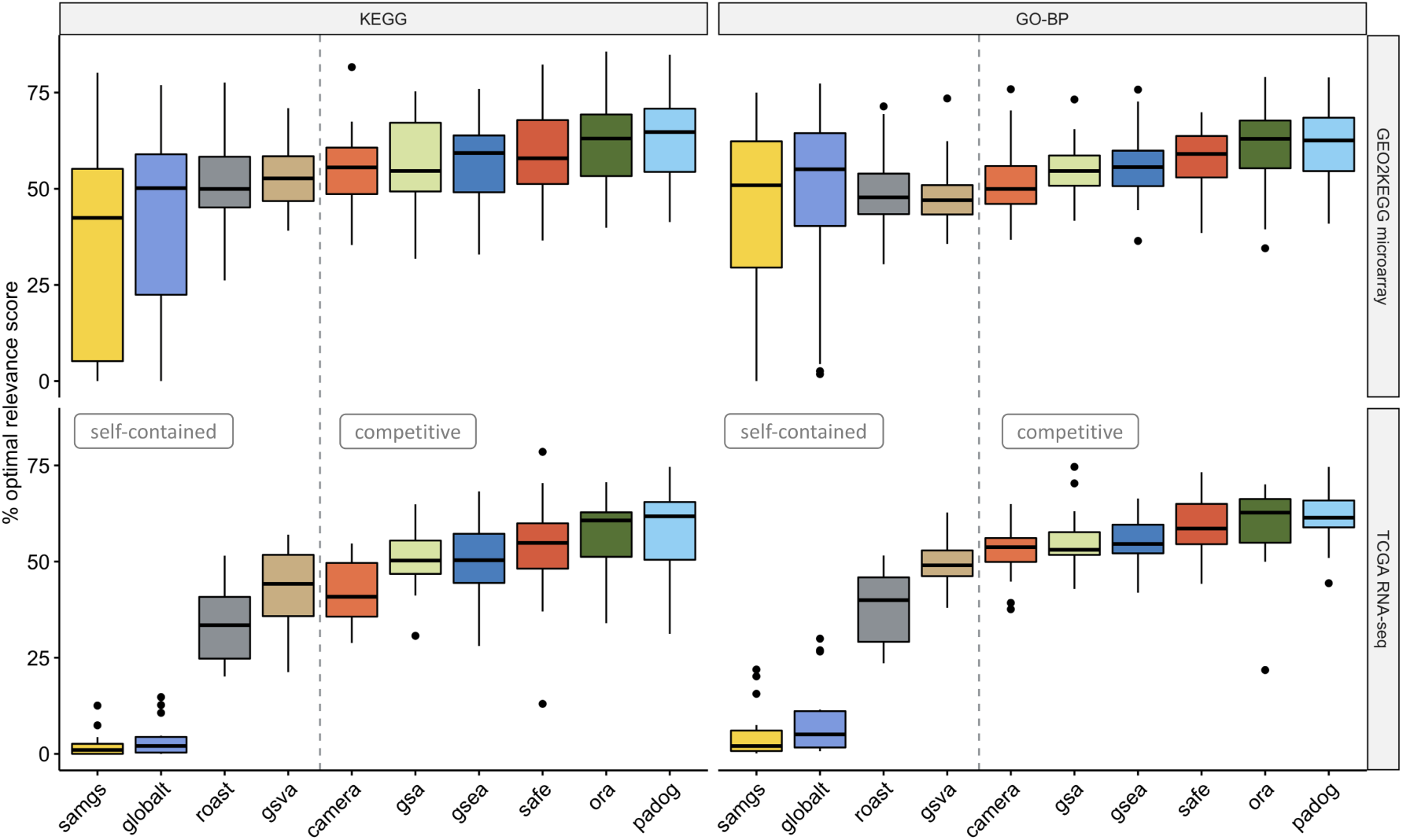
Phenotype relevance. Percentage of the optimal phenotype relevance score (*y*-axis) when applying methods to the GEO2KEGG microarray compendium (top, 42 datasets) and the TCGA RNA-seq compendium (bottom, 15 datasets). Gene sets were defined according to KEGG (left, 323 gene sets) and GO-BP (right, 4,631 gene sets). The grey dashed line divides methods based on the type of null hypothesis tested [4]. Computation of the phenotype relevance score is outlined in Figure 1 and detailed in Methods, Section *Phenotype relevance*.

## Discussion

Gene set enrichment analysis is the mainstay for studying differential regulation of biological processes in high-throughput gene expression data. However, the lack of standards for the assessment of enrichment methods currently prevents to derive reliable conclusions on how existing methods perform relative to each other. It also impairs evaluation of whether newly developed methods provide robust performance improvements over already existing methods.

In this work, we make two important contributions: (1) we implement a framework for standardized and structured benchmarking of EA methods based on defined criteria for applicability, gene set prioritization, and detection of relevant processes. This framework guarantees transparency, reproducibility, and allows straightforward application to additional methods; (2) we perform a systematic assessment of 10 major EA methods, and identified best performing methods, but also general shortcomings in the significance estimation of gene set tests.

In contrast to previous assessments in simulated setups and in the context of microarray data, our benchmark framework encompasses comprehensive collections of curated and fully processed microarray and RNA-seq datasets. These datasets cover a representative range of sample size and differential expression, and allow direct application of EA methods, minimizing differences between assessments that are caused by different preprocessing steps. Complementary to this setup, benchmarking was carried out across the two frequently used KEGG and GO-BP gene set collections, for which the number of gene sets contained and gene set size distribution differ significantly.

To assess the applicability of EA methods, we analyzed (i)computational runtime, and (ii) how methods originally developed for microarray can be applied to RNA-seq data. Runtime evaluation demonstrated moderate differences in applicability that mainly depend on methodological aspects and implementation. Consequently, we found simple parametric tests (CAMERA, ORA, GLOBALTEST) to complete a routine enrichment analysis within seconds on a standard workstation, as compared to computationally more intensive permutation methods (GSA, PADOG, GSEA) that require several minutes. With all evaluated methods thus being in an acceptable range, they provide reference runtimes to compare against when developing new methods. For RNA-seq data provided as raw read counts or TPMs, we found a variance-stabilizing transformation to effectively enable direct applicability of EA methods as for microarray data without notable loss in precision/outcome. Integration of DE RNA-seq tools into EA permutation methods resulted in substantially increased runtimes, and is therefore not recommended without further optimization.

The statistical validity of gene set tests has been repeatedly debated. For example, the independence assumption between genes in ORA or pre-ranked GSEA has been a subject of prolonged controversy [4, 41–44]. On the other hand, the permutation procedure incorporated in many gene set tests has been shown to be biased [45], and inaccurate if permutation *p*-values are reported as zero [46]. Recent studies also reported non-uniform *p*-value distribution that are either systematically biased towards zero (false positive inflation) or one (false negative inflation) [47, 48]. These shortcomings can lead to inappropriately small or large fractions of significant gene sets, and can considerably impair prioritization of gene sets in practice.

Our results demonstrate that the fraction of significant gene sets strongly depends on whether a self-contained or a competitive null hypothesis is tested. Although well-established in theory [4], the practical implications of this distinction are typically not well comprehended. However, we demonstrated that this distinction can in the extreme case determine whether not a single gene set (competitive) or all gene sets (selfcontained) are reported as significant for the same dataset. The drastically different fractions of significant gene sets from competitive and self-contained methods thus require different approaches to interpretation, especially from the perspective of gene set prioritization.

Gene set enrichment analysis is an exploratory rather than confirmatory process, which prompts careful weighing of type I vs. type II error control. For competitive methods, we found the fraction of significant gene sets to be constant across datasets at 5-15% using a nominal significance level of 0.05. This seems to be a fraction of reasonable size for the purpose of generating candidate gene sets for further investigation, when also accepting a certain amount of false positives. Such a compromise is demonstrated by the interesting example of CAMERA, which deliberately abandons strict type I error control by default to compensate for the apparent lack in power of competitive methods [34, 49]. For self-contained methods, we found GSVA and ROAST in conjunction with strict correction for multiple hypothesis testing preferable over other self-contained methods that display a gene set size dependency. However, when stating significance of a self-contained test, we recommend to always report the background level of DE in the dataset, as we found this to determine the rate of significant gene sets. We also note that it is not always trivial to categorize methods as either competitive or self-contained, and that methods combining aspects from both models might either be predominantly or fully competitive or self-contained depending on the execution mode (Supplementary Discussion S2.1).

A critical feature of EA methods is the ability to rank relevant gene sets higher than other gene sets, but a quantitative assessment of phenotype relevance is difficult. Instead of the frequently applied post hoc reasoning on selected enriched gene sets, we *a priori* investigated the association of all gene sets under study with the phenotype. Using the aggregated relevance scores for each gene set, we quantitatively assessed whether certain methods accumulate higher scores towards the top of the ranking. This demonstrated that competitive methods tend to rank relevant gene sets systematically higher than self-contained methods. PADOG consistently returned highest relevance sores, which consolidates and extends previous assessments on microarray data using a single target KEGG pathway per dataset [16, 17].

Although PADOG accumulated higher relevance scores than GSEA, we found ORA to provide equivalent relevance levels as PADOG. This underpins the usefulness of ORA as a fast and effective enrichment method, which might also explain its unbroken popularity [50, 51] despite methodological criticism [4, 41]. However, extrapolation to other ORA implementations should be done with care, as results can differ depending on which genes are considered as DE, and which genes are chosen as the background (Supplementary Discussion S2.2, [6, 49]).

In the absence of a perfect gold standard with established ground truth, our evaluation of phenotype relevance generalizes human evaluation through biological reasoning based on associations reported in the literature. The evaluation thereby remains approximative, and further extension is warranted. This includes (i) replication of our findings on datasets not predominantly focusing on cancer types, and (ii) to resolve cases where the relation between dataset and pre-defined relevant gene sets is not clear-cut. Such an extension to additional datasets and more fine-grained relevance rankings is straightforward in our benchmarking framework and will provide further important steps towards a gold standard for benchmarking of methods for gene set enrichment analysis.

## Methods

### Construction of the benchmark compendia

As illustrated in Figure 1, the two pre-defined benchmark compendia consists of 42 microarray datasets collected by Tarca *et al*. [16, 17] and 33 RNA-seq datasets from The Cancer Genome Atlas [24, TCGA]. These datasets investigate 42 human diseases, including 35 cancer types (Supplementary Tables S1 and S2). Gene set relevance rankings for each disease were constructed by querying the `MalaCards` database [38]. `MalaCards` scores genes for disease relevance based on experimental evidence and co-citation in the literature. Per-gene relevance was summarized across GO and KEGG gene sets by subjecting disease-relevant genes for each disease to `GeneAnalytics` [52].

### Enrichment methods

Enrichment methods selected for assessment are listed in Table 1. Methods were carried out as implemented in the `EnrichmentBrowser` package [20]. Permutation-based methods originally developed for microarray data were assessed in two different ways on RNA-seq data (see column *RNA-seq* in Table 1). As the indicated methods compute *t*-like statistics for each gene in each permutation of the sample labels, we (i) carried these methods out after applying a variance-stabilizing transformation, or (ii) adapted methods to employ RNA-seq specific tools for computation of the per-gene differential expression (DE) statistic in each permutation.

For the variance-stabilizing transformation we used the `cpm` function implemented in the `edgeR` pack-age [27] to compute moderated log2 read counts. Using `edgeR`’s estimate of the common dispersion *ϕ*, the `prior.count` parameter of the `cpm` function was chosen as 0.5 / *ϕ* as previously suggested [33, 53]. On the other hand, methods were adapted as previously described [20] to use `limma`/`voom` [26, 54], `edgeR`, or `DESeq2` [28] for computation of the per-gene statistic in each permutation of sample labels.

### Gene set collections

Human gene set collections were defined according to the GO-BP ontology and the KEGG pathway annotation using the function `getGenesets` from the `EnrichmentBrowser` package. Collections were restricted to gene sets with a minimum and maximum size of 5 and 500, respectively. This yielded 323 KEGG gene sets and 4,631 GO-BP gene sets with a median gene set size of 72 and 11, respectively.

### Runtime

Elapsed runtime was evaluated using the `R` function `system.time` on an Intel Xeon 2.7 GHz machine.

### Statistical significance

The fraction of statistically significant gene sets returned by an EA method when applied to a specific dataset was evaluated with and without multiple testing correction. A nominal significance level of 0.05 was used when not correcting for multiple testing. Multiple testing correction was carried out using the method from Benjamini and Hochberg [25] with an FDR cutoff of 0.05.

*Type I error rate* was evaluated by randomization of the sample labels on the Golub dataset [37]. The dataset contains microarray measurements of acute myeloid leukemia (AML) and acute lymphoblastic leukemia (ALL) patients and is available from Bioconductor in the `golubEsets` data package [55]. Probe level measurements were normalized using the `vsn2` function of the `vsn` package [56]. Normalized data was summarized to gene level using the `probe2gene` function of the `EnrichmentBrowser` package. The type I error rate was estimated for each enrichment method by shuffling the sample labels (ALL vs. AML) 1,000 times and assessing in each permutation the fraction of gene sets with *p* < 0.05.

*Random gene sets* of increasing set size were analyzed to assess whether enrichment methods are affected by gene set size. We therefore sampled 100 random gene sets of defined size *s* 5, 10, 25, 50, 100, 250, 500 and assessed the fraction of significant gene sets for each enrichment method when applied to the Golub dataset using the true sample labels.

### Phenotype relevance

To evaluate the phenotype relevance of a gene set ranking *R*_*m*(*d*)_ obtained from the application of an EA method *m* to an expression dataset *d* investigating phenotype *p*, we assess whether the ranking accumulates phenotype-relevant gene sets at the top. Therefore, we first transform the ranks from the enrichment analysis to weights - where the greater the weight of a gene set, the more it is ranked towards the top of *R*_*m*(*d*)._

#### Transformation of gene set ranks into weights

EA methods return gene sets ranked according to a ranking statistic *S*, typically the gene set *p*-value or gene set score. If the number of gene sets investigated is *N*_*GS*_, then absolute ranks *r*_*A*_ run from 1 to *N*_*GS*_. Relative ranks

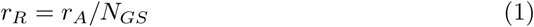

can then be transformed into weights *w* ∈ [0, 1] by

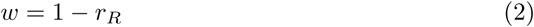

Intuitively, *w* approaches 1 the more a gene set is ranked towards the top of the ranking. In the presence of ties, we calculate relative ranks

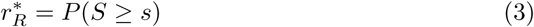

as the fraction of gene sets with a value of the ranking statistic at least as extreme as observed for the gene set to be ranked [20]. Note that 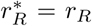 no ties present in the ranking.

#### Relevance score of an EA ranking

To assess the similarity of *R*_*m*(*d*)_ with the corresponding relevance ranking *R*_*p*_ for phenotype *p*, we compute the relevance score

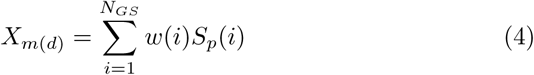

where *w*(*i*) is the weight of gene set *i* in *R*_*m*(*d*)_, and *S*_*p*_(*i*) is the relevance score of gene set *i* in *R*_*p*_. Intuitively, the greater the relevance score *S*_*p*_ of a gene set, the more it is considered relevant for phenotype *p*. Also, the greater the relevance score *X*_*m*(*d*)_ accumulated across the EA ranking, the more similar is the EA ranking *R*_*m*(*d*)_ with the corresponding relevance ranking *R*_*p*_.

#### Random relevance score distribution

To assess the significance of the observed relevance score *X*_*m*(*d*)_, i.e. to assess how likely it is to observe a relevance score equal or greater than the one obtained, we analogously compute relevance scores for random gene set rankings and calculate the *p*-value as for a permutation test.

#### Theoretical optimum

The observed relevance score *X*_*m*(*d*)_ can be used to compare phenotype relevance of two or more EA methods when applied to one particular dataset. However, as the number of gene sets in the relevance rankings can differ between phenotypes, comparison between datasets is not straightforward as resulting relevance scores might scale differently (Supplementary Figures S11 and S12). Therefore, we compute the theoretically optimal score *O*_*p*_ for the case *R*_*m*(*d*)_ = *R*_*p*_ in which the EA ranking is identical to the relevance score ranking. The ratio

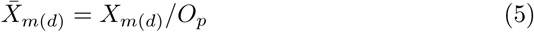

between observed and optimal score can then be used when comparing scores obtained for several methods applied across multiple datasets. This allows to assess whether certain EA methods tend to produce rankings of higher phenotype relevance than other methods when applied to a compendium of datasets.

### Executable benchmark system

The `GSEABenchmarkeR` package is implemented in `R` [57] and is available from `Bioconductor` [58] under http://bioconductor.org/packages/GSEABenchmarkeR. The package allows to (i) load specific pre-defined and user-defined data compendia, (ii) carry out DE analysis across datasets, (iii) apply EA methods to multiple datasets, and (iv) benchmark results with respect to the chosen criteria.

*Loading of benchmark compendia* is facilitated through the `loadEData` function, which simplifies access to (i) the pre-defined GEO2KEGG microarray compendium, (ii)the pre-defined TCGA RNA-seq compendium, and (iii) user-defined data from file. Datasets of the GEO2KEGG microarray compendium are loaded from the Bioconductor packages `KEGGdzPathwaysGEO` and `KEGGandMetacoreDzPathwaysGEO` [16, 17]. Probe-to-gene mapping for each dataset can optionally be carried out, in order to summarize expression levels for probes annotated to the same gene. Datasets of the TCGA RNA-seq compendium are loaded using the `curatedTCGAData` package [59] (TPMs) or from `GSE62944` [60, 61] (raw read counts). User-defined data is also accepted, requiring a file path to the directory where datasets have been saved as serialized `R` data files (Supplementary Methods S1.3).

*Caching* to flexibly save and restore an already processed expression data compendium is incorporated by building on functionality of the `BiocFileCache` pack-age [62]. This is particularly beneficial as preparing an expression data compendium for benchmarking of EA methods can be time-consuming and can involve several pre-processing steps.

*DE analysis* between sample groups for selected datasets of a compendium can be carried out using the function `runDE`. The function invokes `deAna` on each dataset, which contrasts the sample groups depending on data type and user choice via `limma`/`voom`, `edgeR`, or `DESeq2`.

*Enrichment analysis* At the core of applying a specific EA method to a single dataset is the `runEA` function, which delegates execution of the chosen method to either `sbea` (set-based enrichment analysis) or `nbea` (network-based enrichment analysis). Both functions also accept user-defined enrichment methods (Supplementary Methods S1.2, [20]). In addition, `runEA` returns CPU time used and allows saving results for subsequent assessment.

*Parallel computation* of functions for microarray pre-processing, DE analysis, and enrichment analysis when applied to multiple datasets is realized by building on infrastructure implemented in the `BiocParallel` package [63]. Internally, these functions call `bplapply`, which per default triggers parallel computation as con-figured in `BiocParallel`’s registry of computation parameters. As a result, parallel computation is implicitly incorporated when calling these functions on a multi-core machine. To change the execution mode of these functions, accordingly configured computation parameters can either directly be registered, or supplied as an argument to the respective function. Distributed computation on an institutional computer cluster or a computing cloud is straightforward by similarly configuring a computation parameter of class `BatchtoolsParam` for that purpose.

*Benchmarking* Once methods have been applied to a chosen benchmark compendium, they can be subjected to a comparative assessment using dedicated functions for loading, evaluation, and visualization of the results. The function `evalNrSigSets` evaluates the fraction of significant gene sets given a significance level `alpha` and a method for multiple testing correction, which can be chosen from the methods implemented in `p.adjust` from the `stats` pack-age. The function `evalRelevance` evaluates phenotype relevance between EA rankings and corresponding relevance rankings, given a mapping from dataset to phenotype investigated. Integrated relevance rankings can be refined and relevance rankings for additional datasets can also be incorporated (Supplementary Methods S1.4). Detailed documentation of all implemented functions is available in the reference manual of the package.

### Research reproducibility

Results are reproducible using `R` and `Bioconductor`. Code is available from `GitHub` (https://github.com/waldronlab/GSEABenchmarking). The analysis vignette containing literate programming output is available in Additional file 2.

## Acknowledgements

LG was supported by a research fellowship from the German Research Foundation (DFG, GE3023/1-1). LW was supported by grant U24CA18099 from the National Cancer Institute of the National Institutes of Health.

## Additional Files

Additional file 1 — Supplementary Material

PDF document containing Supplementary Figures S1–S14 and

Supplementary Tables S1–S4.

Additional file 2 — Analysis vignette

HTML document containing literate analysis output.

