## Supplementary Material for "Towards a gold standard for benchmarking gene set enrichment analysis"

### Supporting Information

### S1 Supplementary Methods

#### S1.1 Benchmarking network-based methods

Benchmarking with the `GSEABenchmarkeR` package extends to *network-based* methods that incorporate known gene regulatory interactions. For demonstration, we execute two network-based methods (SPIA [1] and GGEA [2]) on three datasets of the GEO2KEGG microarray compendium, and compare their runtimes on these datasets.

```
# setup
> library(GSEABenchmarkeR)
> library(EnrichmentBrowser)

# prepare
> geo2kegg <- loadEData("geo2kegg", nr.datasets=3, cache=FALSE)
> geo2kegg <- maPreproc(geo2kegg)
> geo2kegg <- runDE(geo2kegg)

# get KEGG gene sets
> kegg.gs <- getGenesets(org="hsa", db="kegg")

# compile a gene regulatory network from KEGG
> kegg.grn <- compileGRN(org="hsa", db="kegg")

# execute SPIA and GGEA on the three datasets
> res <- runEA(geo2kegg,
                 methods=c("spia", "ggea"),
                 gs=kegg.gs,
                 grn=kegg.grn,
                 save2file=TRUE,
                 out.dir="~/nbea_bench")

# get the runtimes
> rtimes <- readResults(data.dir="~/nbea_bench", data.ids=names(geo2kegg),
                           methods=c("spia", "ggea"), type="runtime")
> rtimes
$spia
 GSE1297 GSE14762 GSE15471
 188.016  187.544  170.474

$ggea
 GSE1297 GSE14762 GSE15471
  58.351   45.705   46.612

# visualize comparative performance
> bpPlot(rtimes, what="runtime")

# pre-defined network-based methods
> EnrichmentBrowser::nbeaMethods()
[1] "ggea"        "spia"        "pathnet"     "degraph"     "ganpa"
[6] "cepa"        "topologygsa" "netgsa"
```

**Listing 1:** Benchmarking network-based methods

#### S1.2 Benchmarking user-defined methods

User-defined enrichment methods can easily be plugged into the benchmarking framework. For demonstration, we define a dummy enrichment method that randomly draws $p$-values from a uniform distribution. We then execute this method on datasets of the GEO2KEGG compendium and inspect the percentage of significant gene sets returned for each dataset.

```
# defining a new enrichment method
> method <- function(se, gs, alpha, perm)
{
  ps <- runif(length(gs))
  names(ps) <- names(gs)
  return(ps)
}

# execute the method on the three datasets
> res <- runEA(geo2kegg,
                methods=method,
                gs=kegg.gs,
                save2file=TRUE,
                out.dir="~/method_bench")

# get the rankings
> ranks <- readResults(data.dir="~/method_bench", data.ids=names(geo2kegg),
                       methods="method", type="ranking")

# evaluate the percentage of significant gene sets
> sig.sets <- evalSigSets(ranks, padj="none", alpha=0.05)
> sig.sets
                  method
GSE1297         4.012346
GSE14762        3.076923
GSE15471        5.538462
```

**Listing 2:** Benchmarking user-defined methods

#### S1.3 Incorporating user-defined benchmark compendia

The benchmarking can be straightforward extended to additional datasets. The `loadEData` function accepts a directory where datasets of class `SummarizedExperiment` [3] are stored as `RDS` files [4].

```
# chosing a data directory from which additional datasets are loaded
> data.dir <- system.file("extdata", package="GSEABenchmarkeR")
> edat.dir <- file.path(data.dir, "myEData")

# loading from the chosen data directory
> edat <- loadEData(edat.dir)
> names(edat)
[1] "GSE42057x" "GSE7305x"

> edat[[1]]
class: SummarizedExperiment
dim: 50 136
metadata(5): experimentData annotation protocolData dataType dataId
assays(1): exprs
rownames(50): 3310 7318 ... 123036 117157
rowData names(0):
colnames(136): GSM1031553 GSM1031554 ... GSM1031683 GSM1031684
colData names(2): Sample GROUP
```

**Listing 3:** Incorporating user-defined benchmark compendia

#### S1.4 Incorporating user-defined relevance rankings

It is also possible to refine the integrated MalaCards relevance rankings or to incorporate relevance rankings for additional datasets. For demonstration, we define an exemplary relevance ranking for 10 gene sets, and evaluate the relevance accumulated by an exemplary EA ranking.

```
# (1) producing an EA ranking
> ea.ranks <- makeExampleData("ea.res")
> ea.ranks <- gsRanking(ea.ranks, signif.only=FALSE)
> ea.ranks
DataFrame with 10 rows and 2 columns
        GENE.SET       PVAL
     <character> <numeric>
1            gs3      0.007
2            gs4      0.009
3            gs9      0.037
4            gs7      0.039
5            gs6      0.041
6            gs5      0.351
7            gs8      0.437
8           gs10      0.558
9            gs1      0.835
10           gs2      0.978

# (2) defining a relevance score ranking
> rel.ranks <- ea.ranks
> rel.ranks[,2] <- round( runif(nrow(ea.ranks), min=1, max=100) )
> colnames(rel.ranks)[2] <- "REL.SCORE"
> rownames(rel.ranks) <- rel.ranks[,"GENE.SET"]
> ind <- order(rel.ranks[,"REL.SCORE"], decreasing=TRUE)
> rel.ranks <- rel.ranks[ind,]
> rel.ranks
DataFrame with 10 rows and 2 columns
          GENE.SET REL.SCORE
       <character> <numeric>
gs10          gs10        88
gs6            gs6        84
gs9            gs9        70
gs5            gs5        70
gs4            gs4        70
gs8            gs8        62
gs7            gs7        57
gs2            gs2        39
gs3            gs3        22
gs1            gs1        17

# (3a) evaluate relevance score
> evalRelevance(ea.ranks, rel.ranks)
[1] 266.9

# (3b) compute optimal score
> compOpt(rel.ranks, ea.ranks[,"GENE.SET"])
[1] 324.3

# (3c) relevance scores of random gene set rankings
> compRand(rel.ranks, ea.ranks[,"GENE.SET"], perm=3)
[1] 270.2 247.0 278.5
```

**Listing 4:** Incorporating user-defined relevance rankings

### S2 Supplementary Discussion

#### S2.1 Self-contained vs. competitive

It is not always trivial to categorize methods as either competitive or self-contained, and several methods combine aspects from both models. For example, GSEA and SAFE are hybrid in the sense that they motivate their test statistic on the basis of a competitive gene-sampling model, but calculate their $p$-value in a self-contained subject-sampling manner [5]. This similarly applies for GSA, which computes a self-contained test statistic and calculate the $p$-value in a self-contained subject-sampling manner, but uses a competitive gene-sampling procedure for restandardization of the observed and permuted values of the test statistic, making GSA effectively competitive. PADOG largely follows the procedure of GSA by using sample permutation and restandardization via gene permutation, but replaces the maxmean statistic of GSA by a weighted mean that also takes into account the occurrence frequency of genes across all gene sets tested. The classification of GSEA further depends on the execution mode: for small sample sizes, GSEA provides an argument to use gene permutation for the $p$-value calculation, making it fully competitive. When using sample permutation, GSEA can also be executed fully self-contained if the Kolmogorov-Smirnov statistic is calculated on the basis of the DE $p$-values for each gene in the gene set, instead of on their ranks [5].

Another interesting example is GSVA, that belongs to the class of single sample EA methods and thus takes a conceptually different approach than all other methods assessed in this paper. The other methods (1) analyze differential expression of individual genes between sample groups, and (2) summarize DE of individual genes across the gene set of investigation. Applied in a comparison of sample groups, GSVA reverses the typical approach by (1) computing gene set enrichment scores for each sample, and (2) testing for differential "expression" of these enrichment scores between sample groups using e.g. `limma`. While the second step is a self-contained significance assessment of the enrichment scores for each gene set, GSVA computes the enrichment score for each sample like GSEA based on the KS-statistic in an "unsupervised" competitive way, i.e. without taking the sample classification into account [6]. Interestingly, we observed this approach to be effectively self-contained and closely resembling results obtained for ROAST, a fully self-contained method. We further note that such distinctions are not necessary for the competitive methods ORA and CAMERA, and the self-contained methods GLOBALTEST, SAMGS, and ROAST.

#### S2.2 ORA

##### S2.2.1 Choosing the DE genes

DE studies typically report a gene as differentially expressed if the corresponding DE $p$-value, corrected for multiple testing, satisfies the chosen significance level. EA methods that work directly on the list of DE genes are then substantially influenced not only by the DE method, but also by the method for multiple testing correction. ORA is inapplicable if there are few genes satisfying the significance threshold, or if almost all genes are DE. We therefore implemented a flexible, context-dependent adjustment procedure to account for such cases by applying multiple testing correction in dependence on the overall DE level in the dataset:

- the correction method from Benjamini and Hochberg (BH) is applied, if it renders $\geq 1\%$ and $\leq 25\%$ of all measured genes as DE,
- the $p$-values are left unadjusted, if the BH correction results in $< 1\%$ DE genes, and
- the more stringent Bonferroni correction is applied, if the BH correction results in $> 25\%$ DE genes.

Note that resulting $p$-values are not further used for assessing the statistical significance of DE genes within or between datasets. They are solely used to determine which genes are included in the analysis with ORA - where the context-dependent correction ensures that the fraction of included genes is roughly in the same order of magnitude across datasets.

##### S2.2.2 Choosing the background

Competitive gene set tests such as ORA compare the genes of the gene set tested against the background of genes not in the set [5]. Although rarely explicitly stated, the background is thus an important parameter [7], especially for the hypergeometric test used by ORA where it determines the size of the population from which genes are drawn [8]. We consider three different options for the population: (i) all genes measured in the microarray or RNA-seq experiment under study, (ii) all genes annotated in the gene set collection under study, and (iii) the intersection of (i) and (ii). While the differences between these three options seem subtle, the impact on the significance estimation of the hypergeometric test can be substantial.
We illustrate the impact of the choice of the background by considering a transcriptomic study, in which 12,671 genes have been tested for differential expression between two sample conditions and 529 genes were found DE. Among the DE genes, 28 are annotated to a specific gene set, which contains in total 170 genes. This setup corresponds to a 2 x 2 contingency table, where the overlap of 28 genes can be assessed based on the hypergeometric distribution. This corresponds to a one-sided version of Fisher's exact test, yielding here a highly significant enrichment.

```
> deTable <- matrix( c(28, 142, 501, 12000),
             nrow = 2,
             dimnames = list(c("DE", "Not.DE"),
                    c("In.gene.set", "Not.in.gene.set")))
> deTable
        In.gene.set Not.in.gene.set
DE               28             501
Not.DE          142           12000

> fisher.test(deTable, alternative = "greater")

  Fisher's Exact Test for Count Data

data:  deTable
p-value = 4.088e-10
alternative hypothesis: true odds ratio is greater than 1
95 percent confidence interval:
 3.226736      Inf
sample estimates:
odds ratio
  4.721744
```

**Listing 5:** Using all genes measured in the microarray or RNA-seq experiment under study

This setup would be realistic if all genes of the universe have equal chance to be drawn. However, due to overlaps between gene sets and missing annotation for other genes, some genes are preferentially drawn, and some genes cannot be drawn at all. To account for missing annotation, we restrict the population to genes annotated in the gene set collection under study. We illustrate this by using the human KEGG gene set collection that contains roughly 8,000 genes. The resulting $p$-value of the hypergeometric test drops by 4 orders of magnitude.

```
> kegg.gs <- EnrichmentBrowser::getGenesets(org="hsa", db="kegg")
> length(unique(unlist(kegg.gs)))
[1] 7852

> deTable[2,2] <- 8000
> fisher.test(deTable, alternative = "greater")
```

```

  Fisher's Exact Test for Count Data

data:  deTable
p-value = 1.207e-06
alternative hypothesis: true odds ratio is greater than 1
95 percent confidence interval:
 2.150785      Inf
sample estimates:
odds ratio
  3.147949
```

**Listing 6:** Using all genes annotated in the gene set collection under study

A similar argument can be made for genes in the gene set collection that are not measured, which is more common for microarray studies than for RNA-seq studies. To account for such genes, we restrict the population to the intersection of measured genes and annotated genes. For the example considered here, we assume the intersection to be 7,000 genes, reducing the $p$-value by another order of magnitude.

```
> deTable[2,2] <- 7000
> fisher.test(deTable, alternative = "greater")$p.value
[1] 1.240368e-05
```

**Listing 7:** Using the intersection of measured genes and annotated genes

[3] Morgan, M., Obenchain, V., Hester, J., Pages, H.: SummarizedExperiment. doi:10.18129/B9.bioc.SummarizedExperiment

[4] R Core Team: Saving and Restoring R Data Files. https://www.rdocumentation.org/packages/base/versions/3.6.0/topics/readRDS

**Table S1: GEO2KEGG microarray compendium.**

| Dataset | Disease | Disease code |
|---|---|---|
| GSE14924_CD4 | Acute myeloid leukemia | LAML |
| GSE14924_CD8 | Acute myeloid leukemia | LAML |
| GSE9476 | Acute myeloid leukemia | LAML |
| GSE1297 | Alzheimer disease | ALZ |
| GSE16759 | Alzheimer disease | ALZ |
| GSE5281_EC | Alzheimer disease | ALZ |
| GSE5281_HIP | Alzheimer disease | ALZ |
| GSE5281_VCX | Alzheimer disease | ALZ |
| GSE24739_G0 | Chronic myeloid leukemia | CML |
| GSE24739_G1 | Chronic myeloid leukemia | CML |
| GSE23878 | Colorectal cancer | CRC |
| GSE4107 | Colorectal cancer | CRC |
| GSE4183 | Colorectal cancer | CRC |
| GSE8671 | Colorectal cancer | CRC |
| GSE9348 | Colorectal cancer | CRC |
| GSE19420 | Diabetes mellitus type 2 | DMND |
| GSE1145 | Dilated cardiomyopathy | DCM |
| GSE3585 | Dilated cardiomyopathy | DCM |
| GSE7305 | Endometrial cancer | UCEC |
| GSE19728 | Glioma | GBM |
| GSE21354 | Glioma | LGG |
| GSE8762 | Huntington disease | HUNT |
| GSE30153 | Lupus erythematosus systemic | LES |
| GSE18842 | Non small cell lung cancer | LUAD |
| GSE19188 | Non small cell lung cancer | LUAD |
| GSE38666_epithelia | Ovarian neoplasms | OV |
| GSE38666_stroma | Ovarian neoplasms | OV |
| GSE15471 | Pancreatic cancer | PAAD |
| GSE16515 | Pancreatic cancer | PAAD |
| GSE22780 | Pancreatic neoplasms | PAAD |
| GSE32676 | Pancreatic cancer | PAAD |
| GSE20153 | Parkinson disease | PARK |
| GSE20164 | Parkinson disease | PARK |
| GSE20291 | Parkinson disease | PARK |
| GSE6956AA | Prostate cancer | PRAD |
| GSE6956C | Prostate cancer | PRAD |
| GSE11906 | Pulmonary disease chronic obstructive | PDCO |
| GSE42057 | Pulmonary disease chronic obstructive | PDCO |
| GSE14762 | Renal cancer | KIRC |
| GSE781 | Renal cancer | KIRC |
| GSE3467 | Thyroid cancer | THCA |
| GSE3678 | Thyroid cancer | THCA |

**Table S2: TCGA disease codes.**

| Disease code | Disease |
|---|---|
| ACC | Adrenocortical carcinoma |
| BLCA | Bladder Urothelial Carcinoma |
| BRCA | Breast invasive carcinoma |
| CESC | Cervical squamous cell carcinoma and endocervical adenocarcinoma |
| CHOL | Cholangiocarcinoma |
| COAD | Colon adenocarcinoma |
| DLBC | Lymphoid Neoplasm Diffuse Large B-cell Lymphoma |
| ESCA | Esophageal carcinoma |
| GBM | Glioblastoma multiforme |
| HNSC | Head and Neck squamous cell carcinoma |
| KICH | Kidney Chromophobe |
| KIRC | Kidney renal clear cell carcinoma |
| KIRP | Kidney renal papillary cell carcinoma |
| LAML | Acute Myeloid Leukemia |
| LGG | Brain Lower Grade Glioma |
| LIHC | Liver hepatocellular carcinoma |
| LUAD | Lung adenocarcinoma |
| LUSC | Lung squamous cell carcinoma |
| MESO | Mesothelioma |
| OV | Ovarian serous cystadenocarcinoma |
| PAAD | Pancreatic adenocarcinoma |
| PCPG | Pheochromocytoma and Paraganglioma |
| PRAD | Prostate adenocarcinoma |
| READ | Rectum adenocarcinoma |
| SARC | Sarcoma |
| SKCM | Skin Cutaneous Melanoma |
| STAD | Stomach adenocarcinoma |
| TGCT | Testicular Germ Cell Tumors |
| THCA | Thyroid carcinoma |
| THYM | Thymoma |
| UCEC | Uterine Corpus Endometrial Carcinoma |
| UCS | Uterine Carcinosarcoma |
| UVM | Uveal Melanoma |

**Table S3: Dealing with RNA-seq data: correlation of log2 fold changes.** Using the log2 fold changes obtained from applying `voom/limma` to the raw read counts available from GSE62944 as a reference, the table shows Pearson correlation with log2 fold changes obtained from applying `limma` subsequent to a variance stabilizing transformation (VST) on the raw read counts (2nd column), or `voom/limma` on TPMs available from `curatedTCGAData` (3rd column), or `limma` on the log2-transformed TPMs.

| Dataset | VST + limma | TPM + voom/limma | log2 TPM + limma |
|---|---|---|---|
| BLCA | 0.971 | 0.993 | 0.966 |
| BRCA | 0.991 | 0.996 | 0.987 |
| COAD | 0.995 | 0.979 | 0.978 |
| HNSC | 0.982 | 0.998 | 0.982 |
| KICH | 0.99 | 0.996 | 0.983 |
| KIRC | 0.992 | 0.997 | 0.988 |
| KIRP | 0.979 | 0.995 | 0.967 |
| LIHC | 0.964 | 0.995 | 0.956 |
| LUAD | 0.993 | 0.993 | 0.985 |
| LUSC | 0.993 | 0.997 | 0.989 |
| PRAD | 0.991 | 0.992 | 0.985 |
| READ | 0.992 | 0.987 | 0.984 |
| STAD | 0.975 | 0.992 | 0.967 |
| THCA | 0.987 | 0.994 | 0.98 |
| UCEC | 0.985 | 0.992 | 0.978 |

**Table S4: Dealing with RNA-seq data: correlation of -log10 DE *p*-values.** See caption of Table S3.

| Dataset | VST + limma | TPM + voom/limma | log2 TPM + limma |
|---|---|---|---|
| BLCA | 0.976 | 0.98 | 0.963 |
| BRCA | 0.987 | 0.986 | 0.96 |
| COAD | 0.989 | 0.954 | 0.921 |
| HNSC | 0.983 | 0.992 | 0.978 |
| KICH | 0.976 | 0.986 | 0.875 |
| KIRC | 0.985 | 0.989 | 0.964 |
| KIRP | 0.964 | 0.975 | 0.924 |
| LIHC | 0.97 | 0.985 | 0.964 |
| LUAD | 0.989 | 0.971 | 0.963 |
| LUSC | 0.986 | 0.992 | 0.975 |
| PRAD | 0.992 | 0.959 | 0.948 |
| READ | 0.984 | 0.97 | 0.964 |
| STAD | 0.982 | 0.978 | 0.951 |
| THCA | 0.979 | 0.984 | 0.929 |
| UCEC | 0.98 | 0.976 | 0.965 |

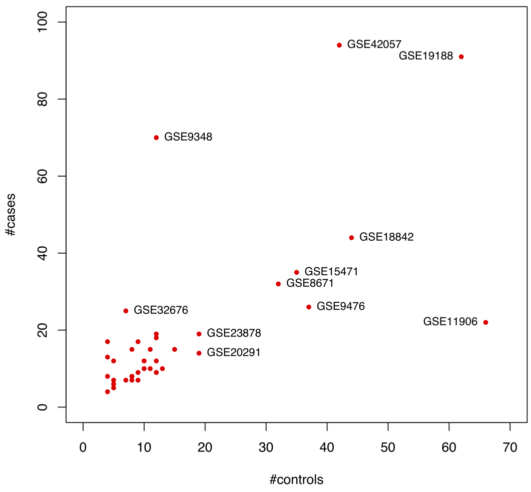

**(a)** GEO2KEGG: number of samples

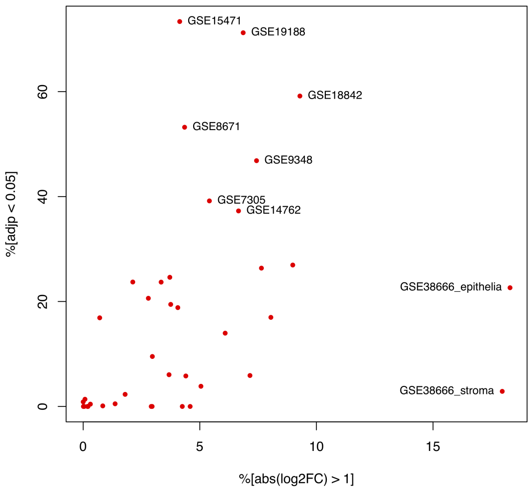

**(b)** GEO2KEGG: percentage of DE genes

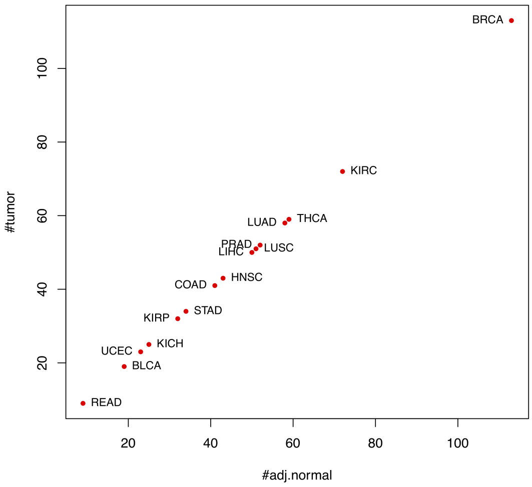

**(c)** TCGA: number of samples

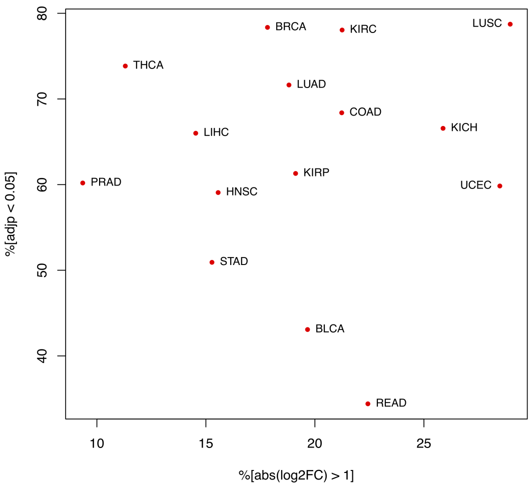

**(d)** TCGA: percentage of DE genes

**Figure S1: Benchmark compendia: sample size and differential expression.** Panel **(a)** and **(c)** show the number of cases and controls for each dataset of the GEO2KEGG microarray compendium ($N = 42$) and the TCGA RNA-seq compendium ($N = 15$), respectively. Using the typical thresholds for differential expression (DE), panel **(b)** and **(d)** show the percentage of genes with an absolute log2 fold change above 1 ($x$-axis) and a Benjamini-Hochberg (BH)-adjusted $p$-value below 0.05 ($y$-axis).

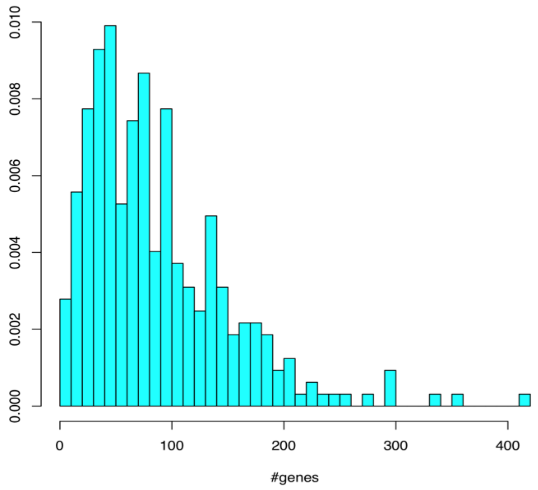

**(a)** KEGG

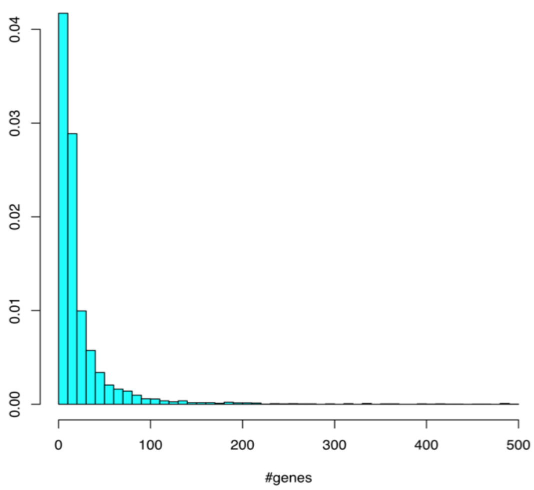

**(b)** GO-BP

**Figure S2: Gene set size distribution.** Considering only gene sets with a minimum of 5 genes and a maximum of 500 genes (the typical thresholds for EA analysis), gene set size distributions are shown for **(a)** 323 human KEGG gene sets (median set size of 72 genes), and **(b)** 4,631 human GO-BP gene sets (median set size of 11 genes). Filtering for set size was applied on a total of 331 KEGG gene sets and 12,078 GO-BP gene sets.

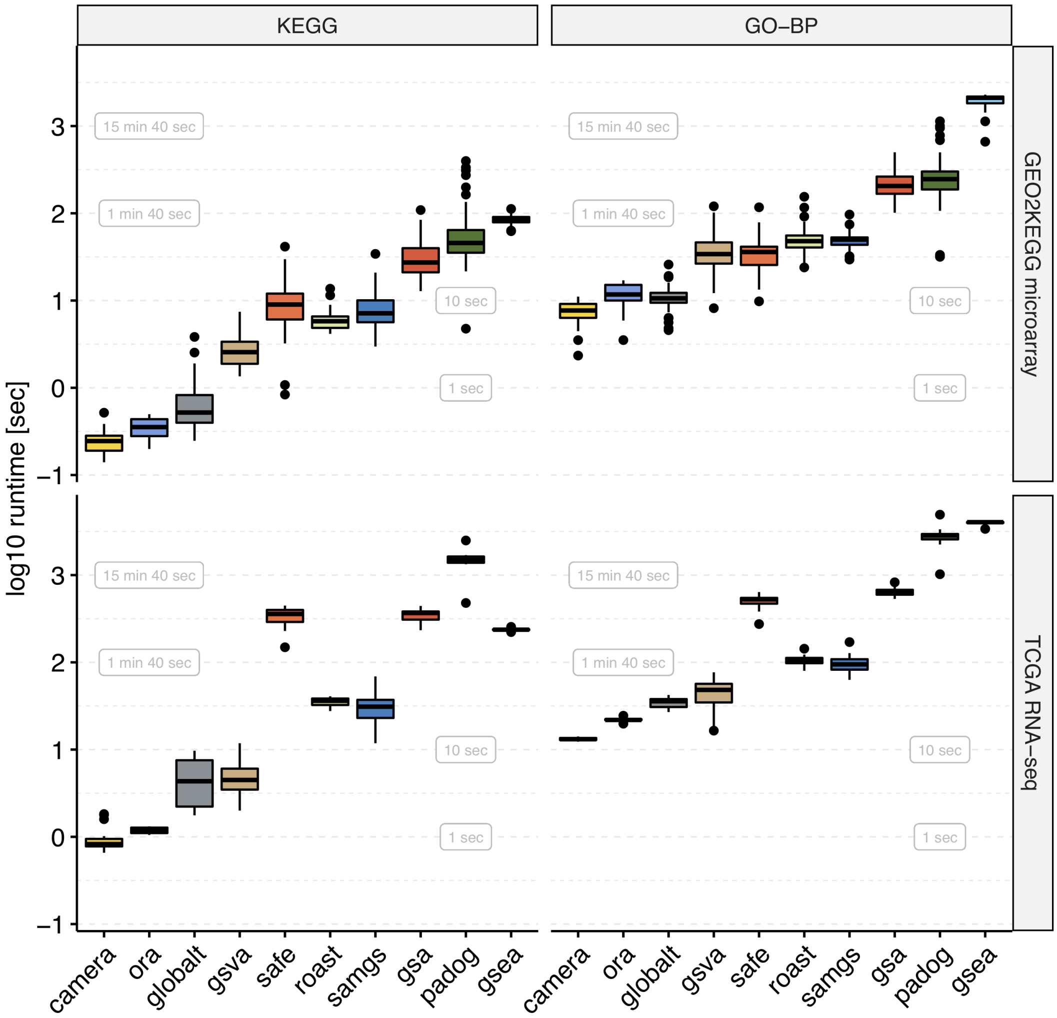

**Figure S3: Runtime.** Shown are the distributions of the elapsed processing times ($y$-axis, log-scale) when applying the enrichment methods indicated on the $x$-axis to the GEO2KEGG microarray compendium (top, 42 datasets) and the TCGA RNA-seq compendium (bottom, 15 datasets). Gene sets were defined according to KEGG (left, 323 gene sets) and GO-BP (right, 4,631 gene sets). Computation was carried out on an Intel Xeon 2.7 GHz machine.

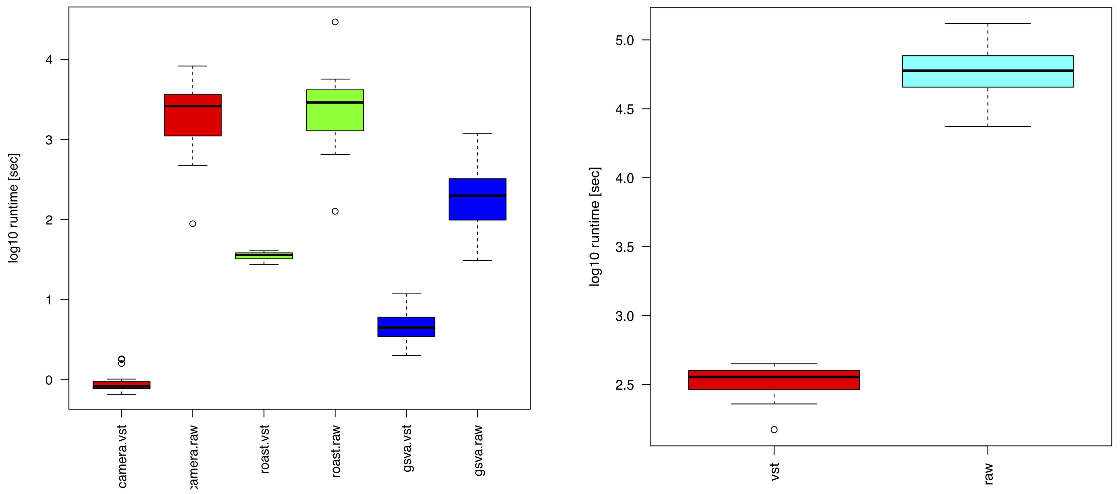

**(a)** CAMERA, ROAST, GSVA

**(b)** SAFE

**Figure S4: Runtime for different RNA-seq modes.** Shown are the distributions of the elapsed processing times ($y$-axis, log-scale) when applying the enrichment methods indicated on the $x$-axis to the TCGA RNA-seq compendium (15 datasets). Gene sets were defined according to KEGG (323 gene sets). Computation was carried out on an Intel Xeon 2.7 GHz machine. VST: application of methods in microarray mode after applying a variance stabilizing transformation (VST) to the raw RNA-seq read counts. RAW: application of methods in RNA-seq mode to the raw RNA-seq read counts. For SAFE, application to the raw RNA-seq read counts was carried out by using `voom`/`limma` for recalculation of the local (per-gene) statistic in each permutation of the sample labels.

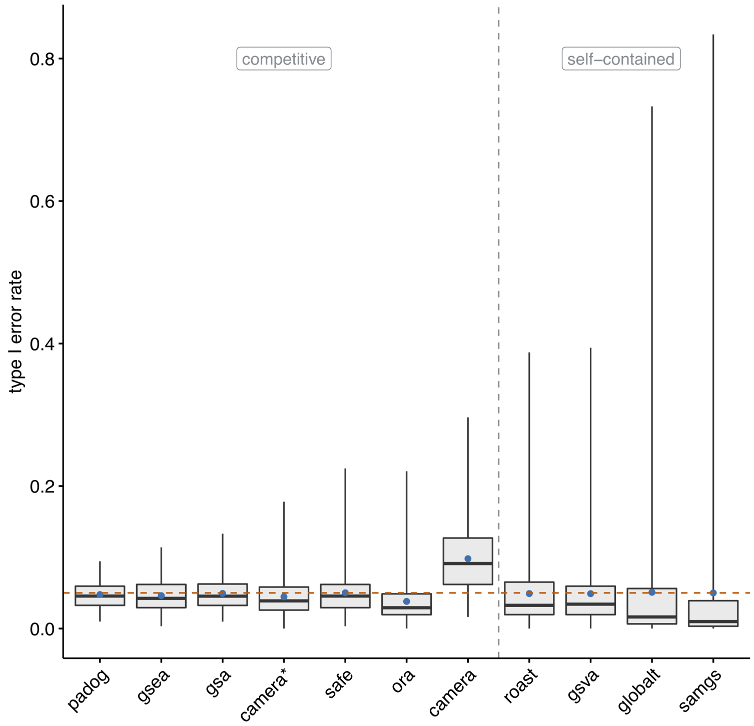

**Figure S5: Random sample labels**. Type I error rates ($y$-axis) as evaluated on the Golub dataset by shuffling sample labels 1000 times, and assessing in each permutation the fraction of gene sets with $p < 0.05$. Gene sets were defined according to KEGG ($N = 323$). Blue points indicate the mean type I error rate. The red dashed line indicates the significance level of 0.05. The grey dashed line divides methods based on the type of null hypothesis tested.

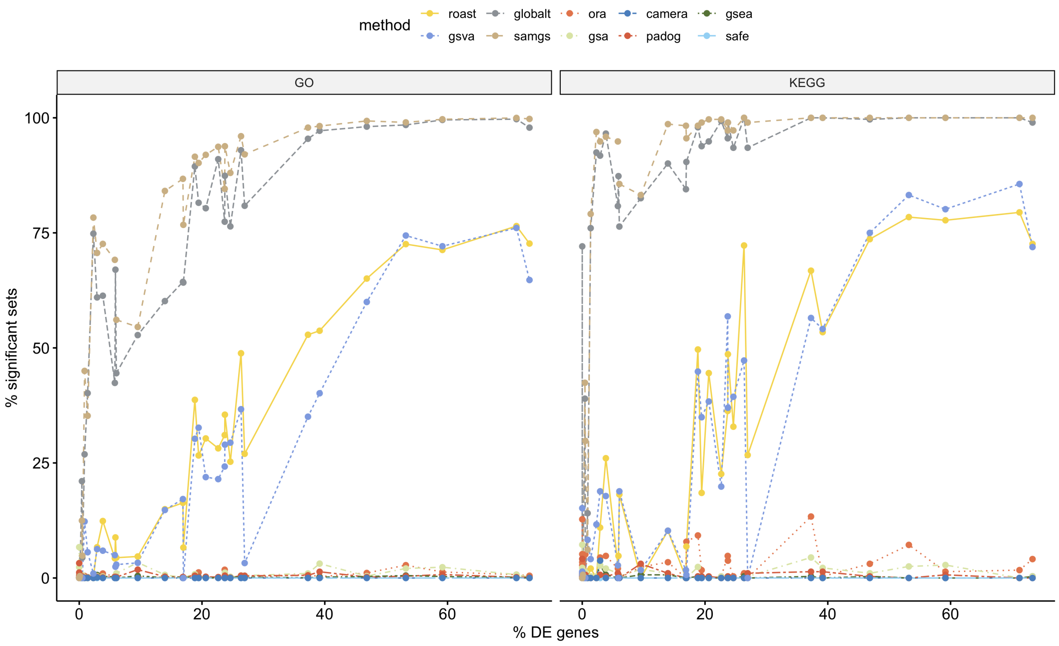

**Figure S6: Correlation of differential expression and gene set enrichment**. Percentage of significant gene sets (FDR $< 0.05$, $y$-axis) when applying methods to the GEO2KEGG microarray compendium (42 datasets) as a function of the percentage of DE genes (FDR $< 0.05$, $x$-axis). Gene sets were defined according to GO-BP (left panel) and KEGG (right panel). The plots show strong correlation for the self-contained methods ROAST (Pearson correlation of 0.968 for GO, and 0.91 for KEGG), GSVA (0.947, 0.903), GLOBALTEST (0.792, 0.637), and SAMGS (0.745, 0.631).

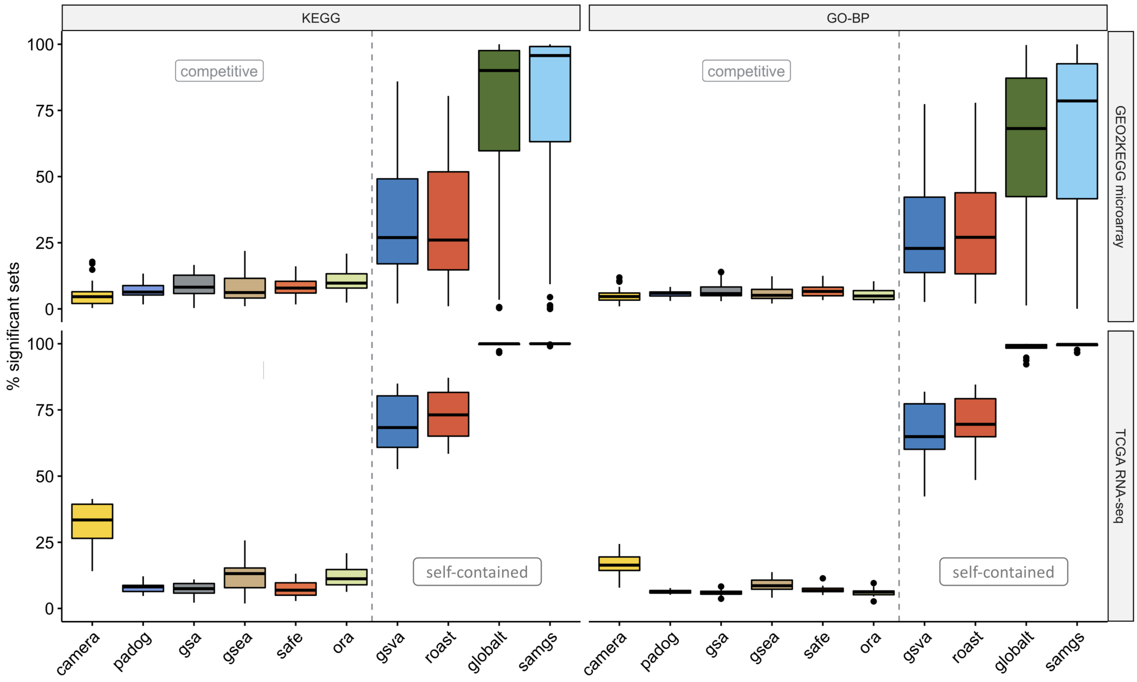

**Figure S7: Statistical significance.** Percentage of significant gene sets before multiple testing correction ($p < 0.05$, $y$-axis) when applying methods to the GEO2KEGG microarray compendium (top, 42 datasets) and the TCGA RNA-seq compendium (bottom, 15 datasets). Gene sets were defined according to KEGG (left, 323 gene sets) and GO-BP (right, 4,631 gene sets). The grey dashed line divides methods based on the type of null hypothesis tested.

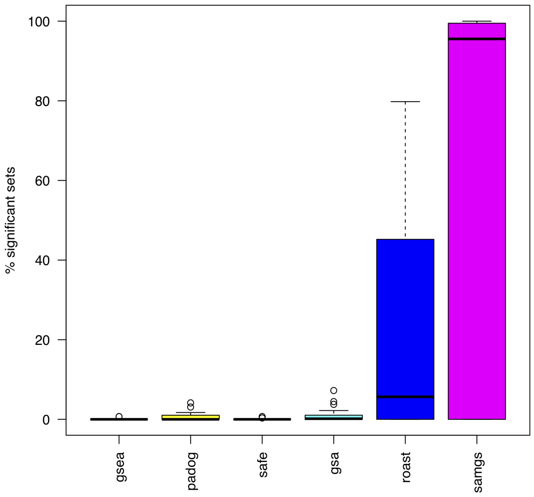

**(a)** p.adjust=BH

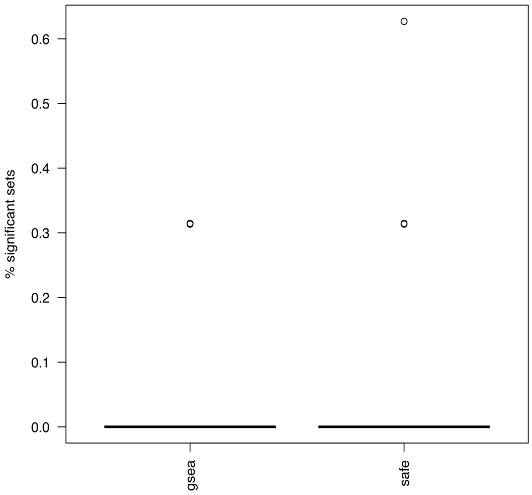

**(b)** built-in FDR

**Figure S8: Statistical significance with 10,000 permutations.** Percentage of significant gene sets (FDR < 0.05, $y$-axis) when applying methods with 10,000 permutations to the GEO2KEGG microarray compendium (42 datasets) and using KEGG gene sets ($N = 323$). Multiple testing correction was carried out with **(a)** the function `p.adjust` from the `stats` package setting the argument `method="BH"`, and **(b)** with the respective built-in FDR correction of GSEA and SAFE.

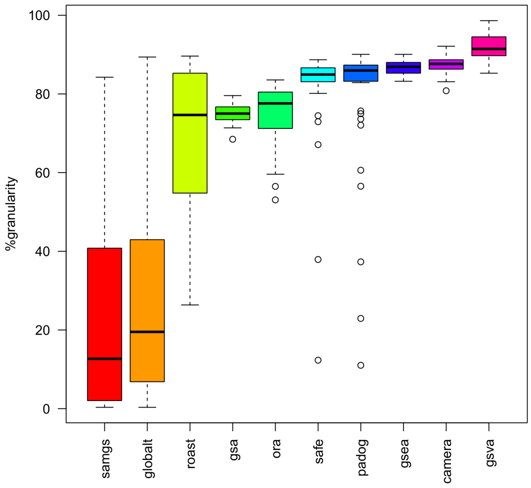

**(a)** KEGG

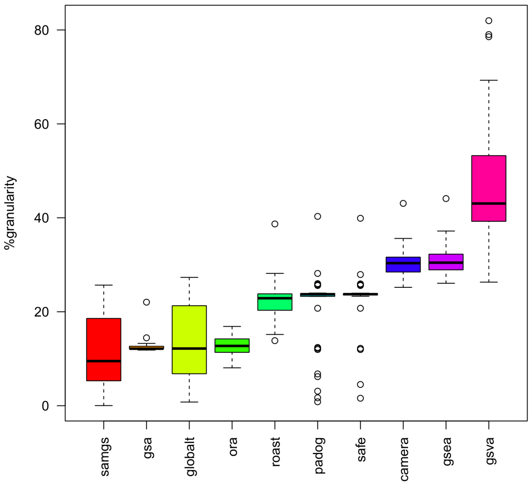

**(b)** GO-BP

**Figure S9: Ranking granularity**. Percentage of gene sets with unique $p$-value returned by SBEA methods when applied to the 42 datasets of the microarray benchmark set and using **(a)** KEGG ($N = 292$), and **(b)** GO-BP ($N = 4128$) gene sets, respectively.

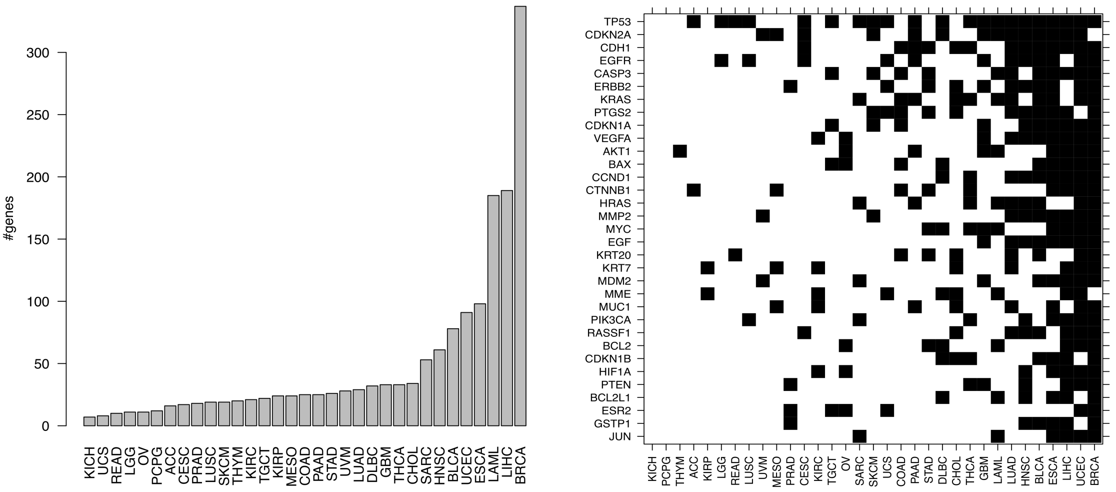

**(a)** Number of genes

**(b)** Most frequent genes

**Figure S10: MalaCards disease genes.**

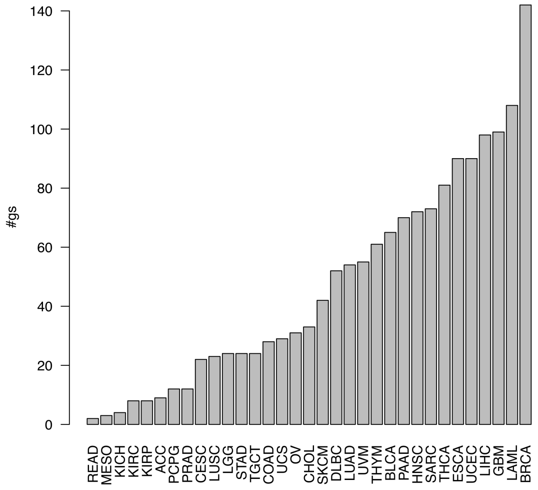

**(a)** KEGG: number of gene sets

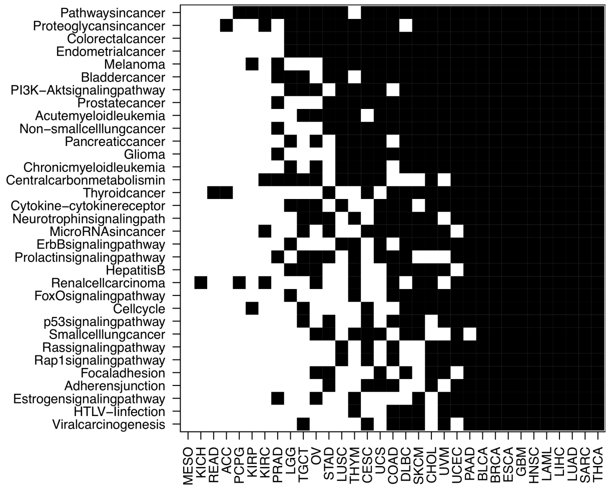

**(b)** KEGG: most frequent gene sets

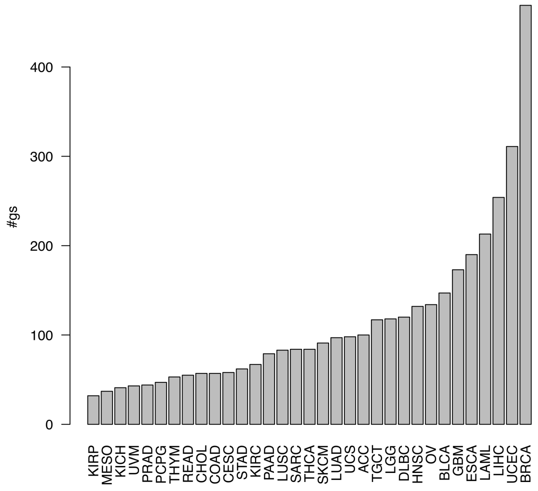

**(c)** GO-BP: number of gene sets

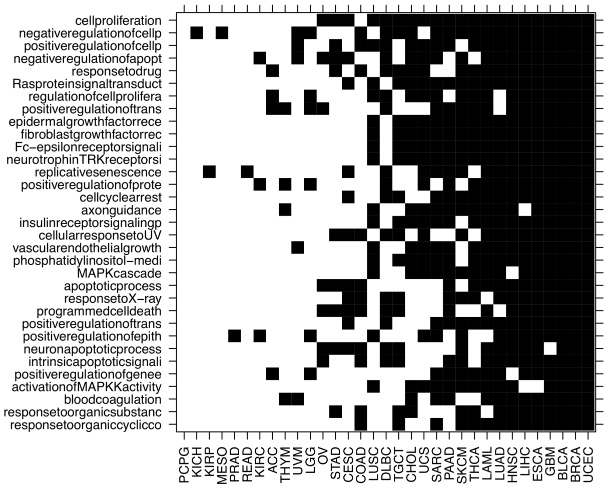

**(d)** GO-BP: most frequent gene sets

**Figure S11: MalaCards disease gene sets.**

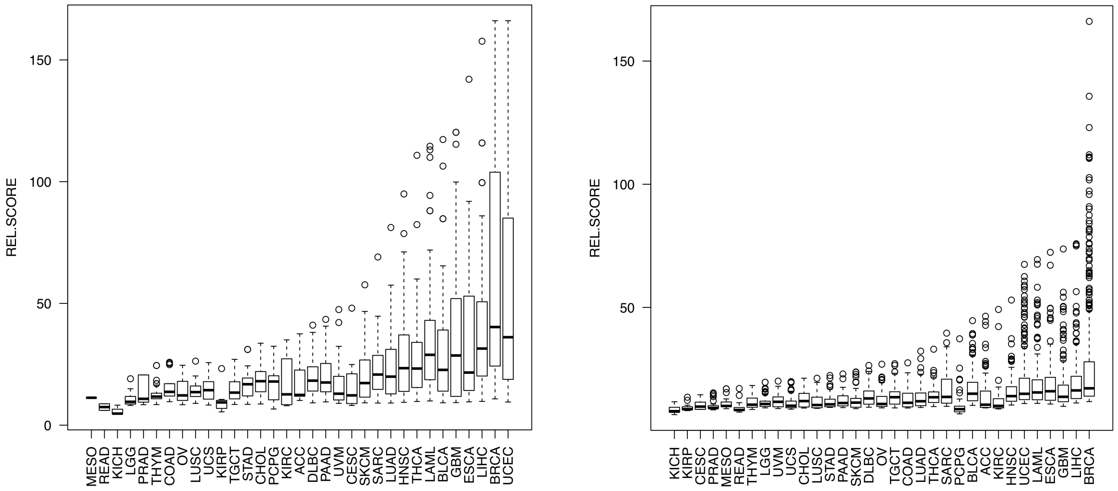

**(a)** KEGG **(b)** GO-BP

**Figure S12: MalaCards relevance score range.**

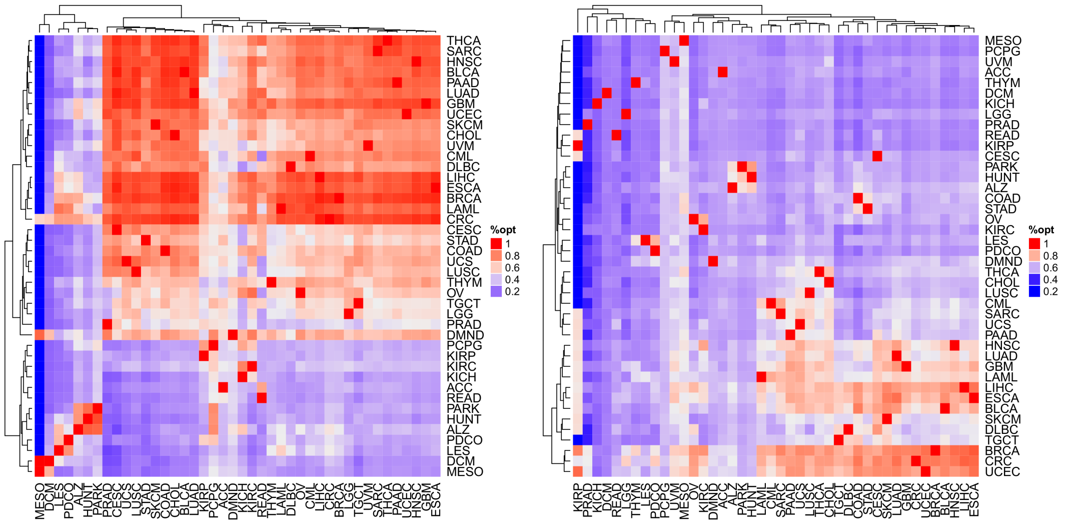

**(a)** KEGG **(b)** GO-BP

**Figure S13: Similarity of MalaCards relevance rankings.** The heatmap shows the percentage of the optimal phenotype relevance score on a color scale. The optimal relevance score corresponds to the case that two relevance rankings are identical. The heatmap in **(a)** for KEGG shows high similarity between the relevance rankings for cancer types (large red cluster in the upper right), neurodegenerative diseases (ALZ, PARK, HUNT), and previously linked autoimmune / chronic inflammatory lung diseases (LES / PDCO, [9]). These similarities are also apparent to a lesser extent for GO-BP relevance rankings in **(b)**, demonstrating higher similarity of relevance rankings within disease classes than between disease classes.

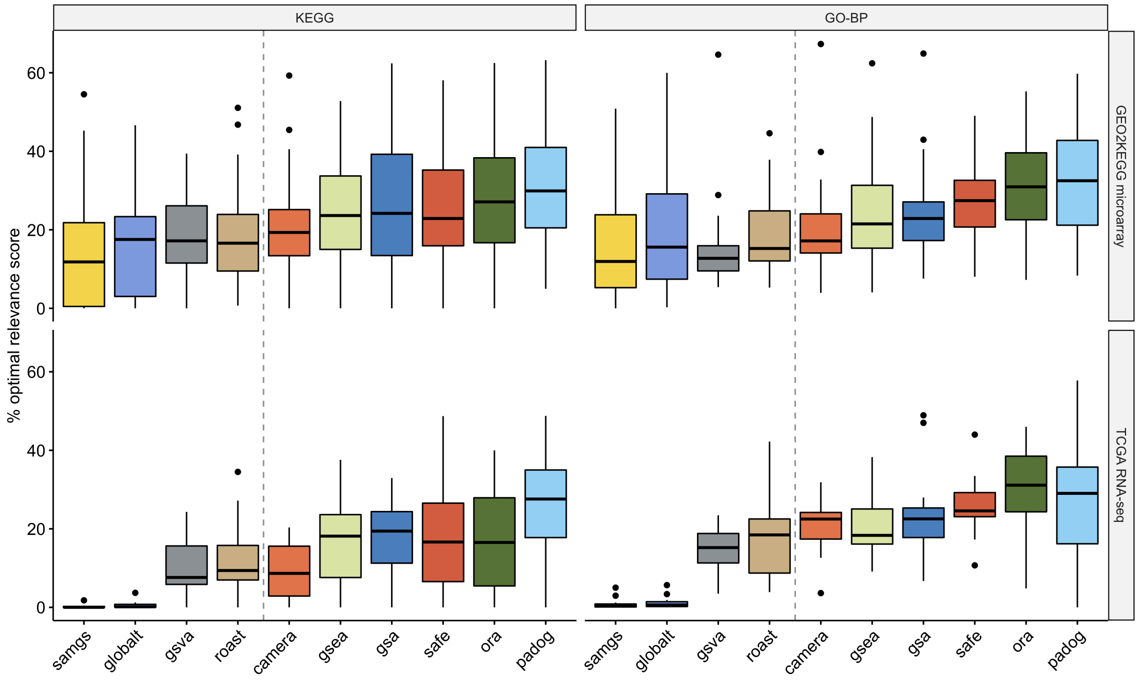

**Figure S14: Phenotype relevance for the top 20% of each EA ranking.** Percentage of the optimal phenotype relevance score ($y$-axis) when applying methods to the GEO2KEGG microarray compendium (top, 42 datasets) and the TCGA RNA-seq compendium (bottom, 15 datasets). Gene sets were defined according to KEGG (left, 323 gene sets) and GO-BP (right, 4,631 gene sets). The grey dashed line divides methods based on the type of null hypothesis tested. Computation of the phenotype relevance score is outlined in Figure 1 of the main manuscript and detailed in Methods, Section *Phenotype relevance*.
